# An Exponential Scale Mixture Model for Metatranscriptomics Data with Application to Inflammatory Bowel Disease

**DOI:** 10.64898/2026.05.15.725552

**Authors:** Hyotae Kim, Li Ma

## Abstract

Metatranscriptomic (MTX) sequencing profiles gene expression across microbial communities, but differential expression analysis is challenging because the abundances are non-negative, extremely sparse, and right-skewed. Standard MTX pipelines output normalized abundances rather than raw counts, limiting the direct use of RNA-seq count-based methods. Gaussian methods require data transformation with pseudocounts, which can lead to distributional misspecification and pseudocount sensitivity. We develop an exponential scale mixture (ESM) framework that directly models MTX abundances with an interpretable mean regression structure while accommodating near-zero concentration and heavy right tails. The complete ESM model, incorporating a two-component mixture and a left-censoring mechanism, can account for extreme sparsity and assign positive probability to observed zeros. In simulation studies, the ESM achieved greater power at comparable empirical false discovery rates (FDRs), more accurate signal strength estimation, better distributional fit, and better FDR calibration than a representative Gaussian alternative. In a case study using inflammatory bowel disease (IBD) multi-omics data, the ESM and the Gaussian model identified markedly different sets of genes (230 vs. 927), while distributional diagnostics strongly favored the ESM. The greater FDR inflation of the Gaussian model in simulations warrants caution in interpreting its larger discovery set. Functional annotation mapped several ESM-exclusive discoveries to gene families associated with IBD-related functions, while Pfam enrichment identified a significant functional signature involving phosphoenolpyruvate (PEP)-utilizing enzymes.

## 1 Introduction

While RNA-seq and scRNA-seq are dominant and well-established sequencing technologies for gene expression analysis, metatranscriptomic sequencing is rapidly gaining traction, as it can profile RNA from entire microbial communities; metatranscriptomic (MTX) analysis thereby enables investigations that connect community genetic potential with molecular activity (e.g., Franzosa et al. 2014, Bashiardes et al. 2016, Lloyd-Price et al. 2019).

To process metatranscriptomic sequencing data, several MTX pipelines—integrated collections of bioinformatics tools—have been developed. HUMAnN (Franzosa et al. 2018, Beghini et al. 2021) is one representative example; a comprehensive comparison of existing pipelines can be found in Shakya et al. (2019). These workflows typically produce transcript abundances normalized by gene length as RPK (reads per kilobase) and then further normalized by sequencing depth, reported as RPKM (reads per kilobase per million mapped reads) or CPM (copies per million), rather than as raw (i.e., unnormalized) counts. Hence, count-based methods originally developed for RNA-seq and scRNA-seq are no longer directly applicable off the shelf. Even when raw counts are available, you still need additional information on the length of all identified genes, and no known optimal normalization or adjustment method exists for gene length within count-based models, which further complicates their use.

Zhang et al. (2021) introduced Gaussian-based modeling methods for processed MTX abundance data that adjust for DNA gene copy number (also referred to as DNA abundance or metagenomic (MGX) abundance), thereby accounting for variation in taxonomic composition and genomic potential across observations that may otherwise confound differential expression analysis. Other methods developed for RNA-seq that do not directly model raw counts may also be adapted to MTX data with adjustment for DNA abundance, including zero-inflated beta regression (Peng et al. 2016) and truncated Gaussian hurdle models (Finak et al. 2015).

We propose a novel modeling framework based on a scale mixture of exponential distributions for differential expression analysis of MTX data. The exponential scale is represented as the product of a mean-determining component and an observation-specific latent scale that captures additional heterogeneity. Under a specific inverse-gamma prior, marginalizing over the latent scale yields a Lomax distribution with an interpretable mean-regression structure and precision controlled by the inverse-gamma hyperparameter, with the regression coefficients representing log fold changes in mean abundance.

A useful property of the base framework is that the precision parameter – the inverse-gamma hyperparameter – modulates both the concentration of probability near zero and the heaviness of the right tail: smaller values place more probability near zero while inducing heavier tails. This feature may be helpful for representing MTX data, which typically exhibit a concentration near zero together with a small number of large nonzero abundances, through the use of a small value of the precision parameter. However, extreme sparsity in the data may lead to a tiny value of the precision parameter, causing the base framework to allocate too much probability to the region near zero and leave very little probability over the remaining positive range. As a result, the model may inadequately represent positive abundances despite the increasingly heavy right tail. To tackle this, our complete model introduces an additional mixture layer with two distinct precision parameters. One component accommodates the strong concentration near zero, while the other allows appreciable density over the positive range. We also propose a left-censoring mechanism to take into account recorded zeros arising from latent abundances below a detection threshold. From a sampling-distribution perspective, this mechanism enables exact zeros to be generated under the model by treating latent abundances below the threshold as zeros.

The rest of the paper is organized as follows. Section 2 introduces the motivating application, with Section 2.1 describing the data generated by the HUMAnN pipeline and Section 2.2 discussing limitations of an existing Gaussian-based approach. Section 3 develops our modeling method, beginning with the base framework and its distributional properties in Sections 3.1 and 3.2, respectively, followed by the complete model in Section 3.3. Section 4 evaluates the proposed approach through simulation studies and comparison with a representative Gaussian-based model, and Section 5 presents the case study. Section 6 concludes the paper.

## 2 Motivating Application

### 2.1 MTX Data Description

In this paper, we analyze the Inflammatory Bowel Disease Multi’omics Database (IBDMDB; Lloyd-Price et al. 2019), available at https://ibdmdb.org. The resource provides both MGX and MTX data processed with the HUMAnN pipeline, along with associated metadata. These include community-level gene family abundances, each partitioned into contributions from individual organisms.

Table 1 presents a subset of the MTX data for the gene family UniRef90_A0A015QIN6. Columns other than the first, which contains feature descriptions, correspond to samples, three of which are shown here and selected arbitrarily. The first row reports the aggregate gene family abundance, equal to the sum of the subsequent species-level abundances (prefixed by s), with the corresponding genus-level information prefixed by g. The last row, labeled “unclassified”, represents reads mapped to the gene family but lacking taxonomic assignment. All entries in the table are nonnegative real values representing CPM-scaled abundances.

**Table 1:**
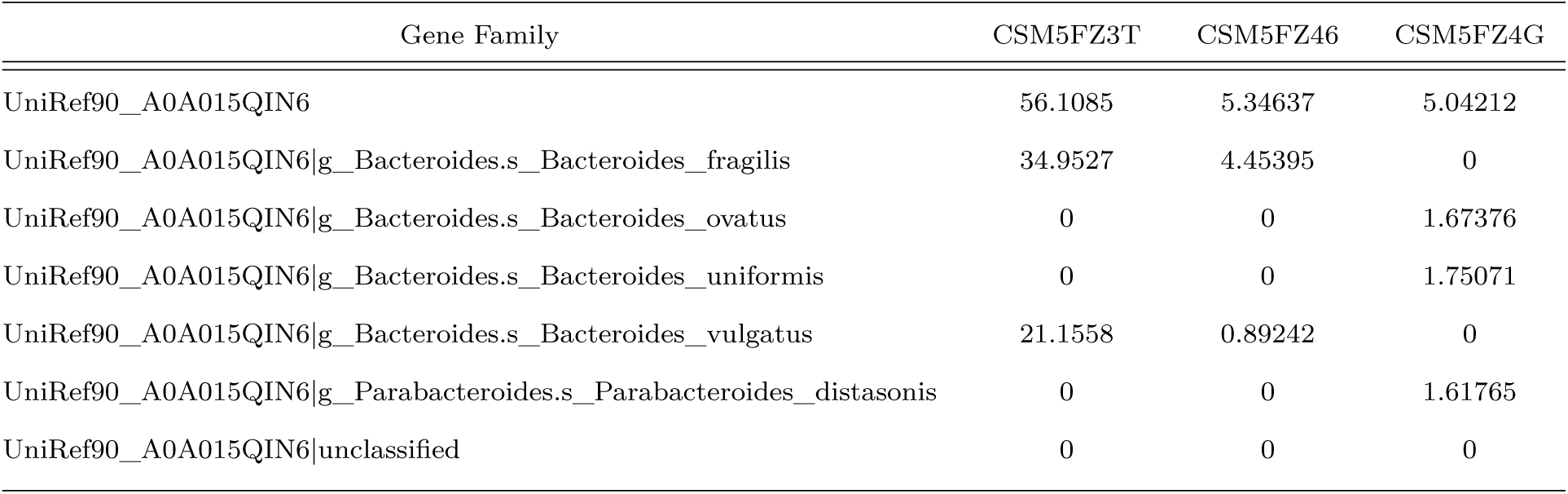
Subset of MTX data showing transcript abundance for one gene family UniRef90_A0A015QIN6 across three samples (columns).

| Gene Family | CSM5FZ3T | CSM5FZ46 | CSM5FZ4G |
|---|---|---|---|
| UniRef90__A0A015QIN6 | 56.1085 | 5.34637 | 5.04212 |
| UniRef90__A0A015QIN6\|g__Bacteroides.s__Bacteroides__fragilis | 34.9527 | 4.45395 | 0 |
| UniRef90__A0A015QIN6\|g__Bacteroides.s__Bacteroides__ovatus | 0 | 0 | 1.67376 |
| UniRef90__A0A015QIN6\|g__Bacteroides.s__Bacteroides__uniformis | 0 | 0 | 1.75071 |
| UniRef90__A0A015QIN6\|g__Bacteroides.s__Bacteroides__vulgatus | 21.1558 | 0.89242 | 0 |
| UniRef90__A0A015QIN6\|g__Parabacteroides.s__Parabacteroides__distasonis | 0 | 0 | 1.61765 |
| UniRef90__A0A015QIN6\|unclassified | 0 | 0 | 0 |

The MTX data consist of 817 samples and 1, 088, 367 gene families, each with associated taxa. For the gene family UniRef90_A0A015QIN6, approximately 55.45% of the species-level observations are zero-valued. The left panel of Figure 1 displays the distribution of the species-level observations for this gene, ranging from 0 to 54.6453, with the 75th percentile at 0.4846, indicating a highly right-skewed distribution.

**Figure 1:**
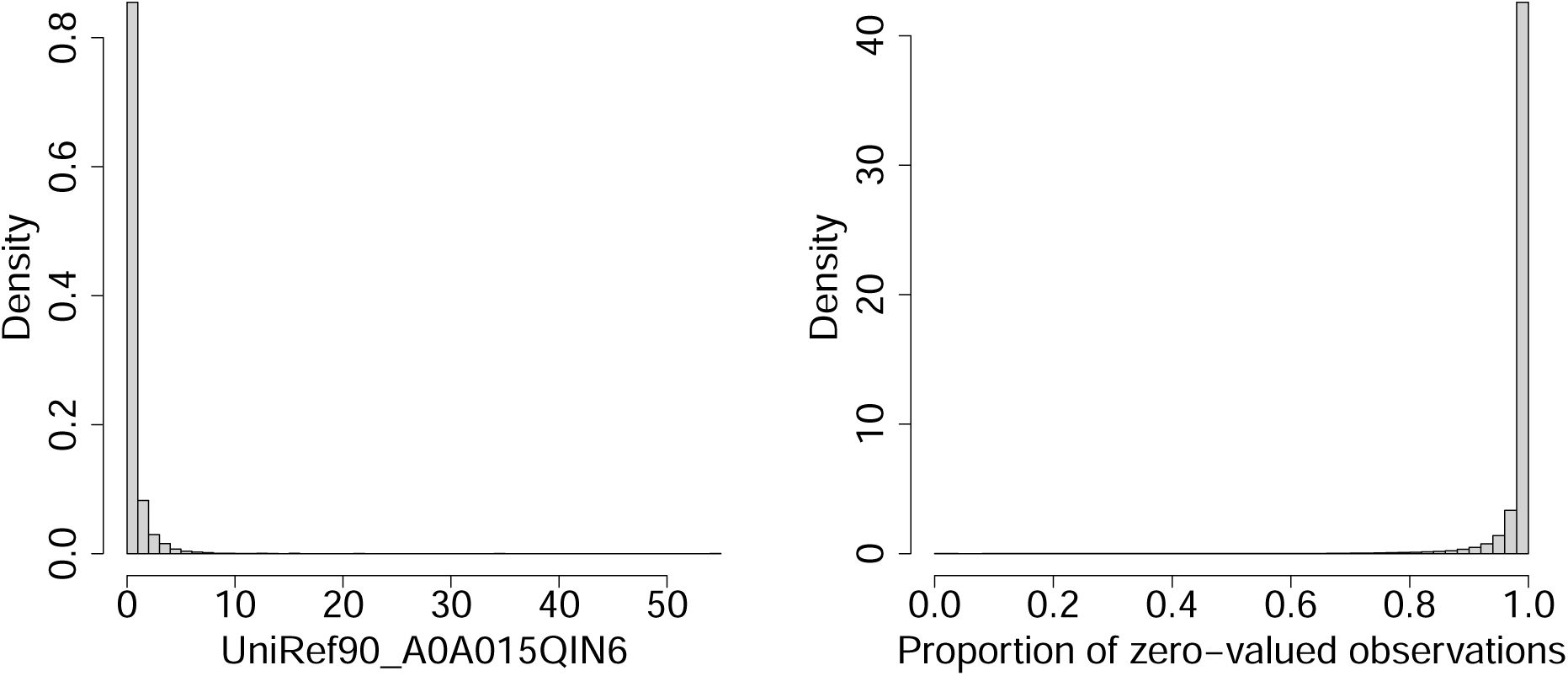
Histogram of transcript abundances for observations in the gene family UniRef90_A0A015QIN6 (left) and histogram of the proportion of zero-valued observations across gene families (right).

The histogram in the right panel presents the distribution of the proportion of zero-valued observations across all gene families. The proportions range from 0 to 0.9988, with mean (median) values of 0.9854 (0.9963). The distribution is highly left-skewed, with even the 25th percentile at 0.9896.

Our approach, introduced in the following section, models the MTX data while incorporating the associated MGX data, provided in the same tabular format from the database, as a covariate for adjustment. As a preliminary step, we construct species-level paired abundance data for each gene family by removing the aggregated (first) and “unclassified” (last) rows and retaining only taxa common to both the MTX and MGX data sets.

Throughout the paper, we use the term “gene” to refer to an annotated gene family (UniRef90 cluster) as reported by the HUMAnN pipeline.

### 2.2 Limitations of Existing Gaussian Approaches

Figure 2 illustrates the limitations of the Gaussian approach. This approach requires a log transformation of the data, which in turn necessitates the addition of a pseudocount. The first four panels show the residual density from the Gaussian model, with log-transformed MGX abundance included as a covariate, under different choices of the pseudocount. One choice (PC4), used in Zhang et al. (2021), sets the pseudocount equal to one-half of the minimum nonzero value in the MTX and MGX data sets, respectively, to replace zero abundances in each data set. As suggested therein, sample pairs with both MTX and MGX equal to zero are excluded before model fitting; we discuss such filtering steps further in Section 3.1.

**Figure 2:**
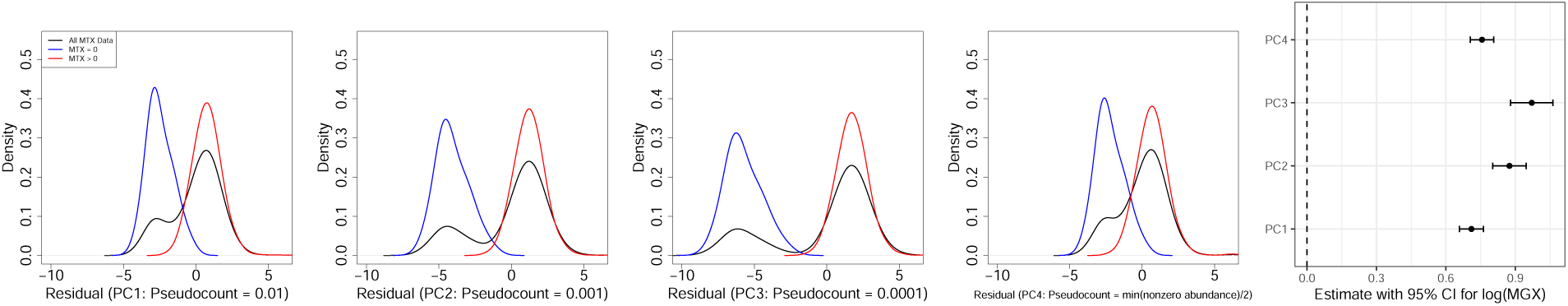
(UniRef90_A0A015QIN6) Density plots (via density() in R) of residuals from the linear model, lm(), under four pseudocount choices (0.01, 0.001, 0.0001, half the minimum nonzero abundance) are shown in the first four panels, followed by the coefficient estimates with 95% confidence intervals for MGX on the log scale.

The resulting residual densities exhibit pronounced bimodality, with the two peaks aligning with the residual densities for positive MTX observations and zero observations, respectively.

The location of the latter varies with the chosen pseudocount. Hence, the unimodal Gaussian specification is inadequate for describing the data. Moreover, the coefficient estimates in the final panel demonstrate that the results also depend on the choice of pseudocount, but no universally accepted optimal value exists. Consequently, inference from Gaussian-based models might be distorted by this choice. The distributional inadequacy and sensitivity to the pseudocount motivate our model development.

## 3 Modeling Approach

We first introduce the base exponential scale mixture framework in Section 3.1 and establish its key distributional and inferential properties in Section 3.2. To address a potential limitation of the base framework under extreme data sparsity, we develop the complete model in Section 3.3 by adding an extra mixture layer with two distinct precision parameters. The complete model also incorporates a left-censoring mechanism for zero-valued measurements.

### 3.1 Base Modeling Framework

For a given gene, let *y_ij_* ∈ R^+^ denote the scaled RNA abundance (e.g., RPKM or CPM) for taxon *j* = 1*, . . ., J* ^∗^ in sample *i* = 1*, . . ., n*^∗^. Although the data can also be indexed by gene as *y_ijs_*, with corresponding dimensions *J_s_* ^∗^ and *n_s_*^∗^, where *s* denotes the gene, we suppress the index *s* for notational simplicity. The following framework is applied independently to each gene, so it can be implemented in parallel across genes.

In designing the model, we consider three key features of MTX abundance data: nonnegative support, substantial concentration at or near zero, and a long right tail with relatively large nonzero observations. The exponential distribution addresses the first two features, as it is supported on the nonnegative real line and has its mode at zero. Moreover, its scale parameter is also its mean, facilitating an interpretable mean-regression structure. However, the exponential distribution is too restrictive to capture the long right tail observed in MTX data. We therefore formulate the exponential scale as the product of a mean component and a latent random scale, leading to an exponential scale mixture formulation. Assigning a particular inverse-gamma distribution to the latent scale and marginalizing over it yields a Lomax distribution that retains the interpretable mean-regression structure while providing a heavier right tail than the exponential distribution. Let Lo(*a, b*) denote the Lomax distribution with shape *a >* 0 and scale *b >* 0, and mean *b/*(*a* − 1) for *a >* 1.

The marginal form of the base framework is

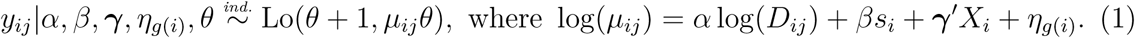

Let Exp(*λ*) denote the exponential distribution with scale parameter *λ*. The marginal model in (1) admits the following equivalent hierarchical representation:

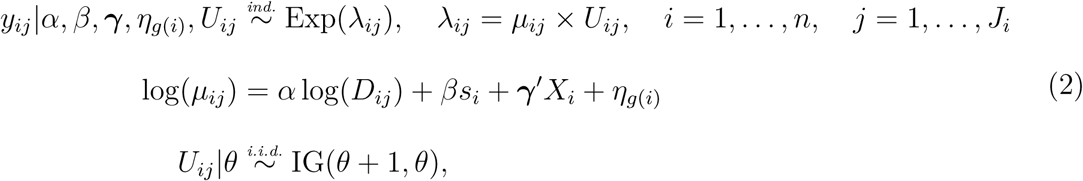

where IG(*a, b*) denotes an inverse-gamma distribution with mean *b/*(*a* − 1) and variance *b*^2^*/*[(*a* − 1)^2^(*a* − 2)] for *a >* 2. *U_ij_* captures additional measurement-level heterogeneity beyond the systematic variation in mean abundance reflected by *µ_ij_*. A key benefit of the hierarchical representation is that it facilitates posterior inference by yielding closed-form full conditional distributions for all model parameters except *θ* (see Section F.2 of the Supplementary Material). It also provides the foundation for the complete model specified in Section 3.3. Section 3.2 establishes the equivalence between the marginal and hierarchical representations in (1) and (2), respectively, and examines several key properties.

*D_ij_* represents the scaled DNA abundance, and *s_i_* is an observed covariate used in differential expression analysis. The vector *X_i_* = (1*, x_i_*_1_*, . . ., x_ip_*)^′^ contains an intercept and *p* additional covariates of interest, with associated coefficients ***γ*** = (*γ*_0_*, . . ., γ_p_*)^′^. In addition, *η_g_*_(_*_i_*_)_ is a subject-specific random effect, where each subject *g*(*i*) may contribute multiple samples in the longitudinal setting. *J_i_* and *n* denote the sample-specific number of taxa and the number of samples, respectively, remaining after the filtering procedure described later, where *J_i_* ≤ *J* ^∗^ and *n* ≤ *n*^∗^. Because the set of taxa may vary across samples after filtering, *J_i_* is sample-specific.

As in Zhang et al. (2021), we incorporate MGX abundance (*D_ij_*) into the regression structure to alleviate the risk of spurious differential expression discoveries, since differences in RNA abundance may reflect differences in gene copy number rather than true differences in expression levels (see Figure 1A of the reference paper for illustrative cases). Accounting for DNA abundance allows for disentangling transcriptional differences from those arising due to variation in gene copy number.

MGX abundance is also used for filtering. Zhang et al. (2021) views the “semi-strict” (pre-)filtering procedure, which removes observations when both MTX and MGX values are zero, as providing a balance between eliminating technical zeros and retaining sufficient observations, and compares it with two alternatives that are more permissive (“lenient”) or more stringent (“strict”). We extend the “semi-strict” filtering by excluding any measurement with zero MGX, even when the paired MTX measurement is positive, because observed transcript abundance in the absence of corresponding detectable DNA abundance is unusual and may indicate technical artifacts.

Lastly, for the analyses in Sections 4 and 5, we use RNA and DNA abundances in CPM for *y_ij_* and *D_ij_*, respectively. The grouping variable for differential expression analysis, *s_i_*, is a binary indicator distinguishing dysbiotic samples, characterized by substantial deviations from non-IBD reference MGX profiles, from non-dysbiotic samples (see Lloyd-Price et al. 2019, for details). We treat consent age and antibiotic use as covariates *x_ik_*, *k* = 1, 2, and participant ID as the subject index *g*(*i*). All data were obtained from the supplementary files *HMP2 Metadata* and *Dysbiosis Scores*, which accompany the MTX-MGX abundance data in the IBDMDB.

### 3.2 Key Properties of the Base Framework

Under the modeling framework in (2), marginalization over the auxiliary variable *U_ij_*with its inverse-gamma prior yields a Lomax distribution in (1) (Proposition A.1, Supplementary Material), with mean *µ_ij_* and variance *µ_ij_*^2^ (*θ* + 1)*/*(*θ* − 1) for *θ >* 1, and infinite variance for 0 *< θ* ≤ 1 (Proposition A.2, Supplementary Material).

The coefficient *γ_k_* for *x_ik_* is interpreted as the log fold change in the mean (as shown in Proposition A.3 of the Supplementary Material); analogous interpretations apply to *α* and *β*. The inverse-gamma hyperparameter *θ*, which modulates the variance, serves as a precision parameter.

Figure 3 displays the probability density and cumulative distribution functions of the Lomax distribution, which is the sampling distribution for *y_ij_* under our base model. One notable characteristic is its support on the nonnegative real line, including zero. As such, the model can be applied directly to zero-valued observations in the likelihood without transformation (i.e., without a pseudocount). Another property worth noting is the fact that *θ* modulates not only the variance but also the concentration of probability near zero as well as the heaviness of the tail. Smaller values of *θ* place more probability near the origin and simultaneously induce heavier tails. Thus, the model can capture substantial concentration near zero and large nonzero measurements. As shown in the right panel, the smallest value of *θ* (*θ* = 0.5; blue curve) yields the smallest 0.9 quantile, reflecting greater concentration near zero. At larger values of *y*, however, its cumulative distribution function increases more slowly than those corresponding to larger values of *θ*, indicating a heavier tail. Formally, the model exhibits a polynomial tail with tail index (*θ* + 1) (Proposition A.4, Supplementary Material), which is heavier than the tail of the log-normal distribution under the Gaussian model.

**Figure 3:**
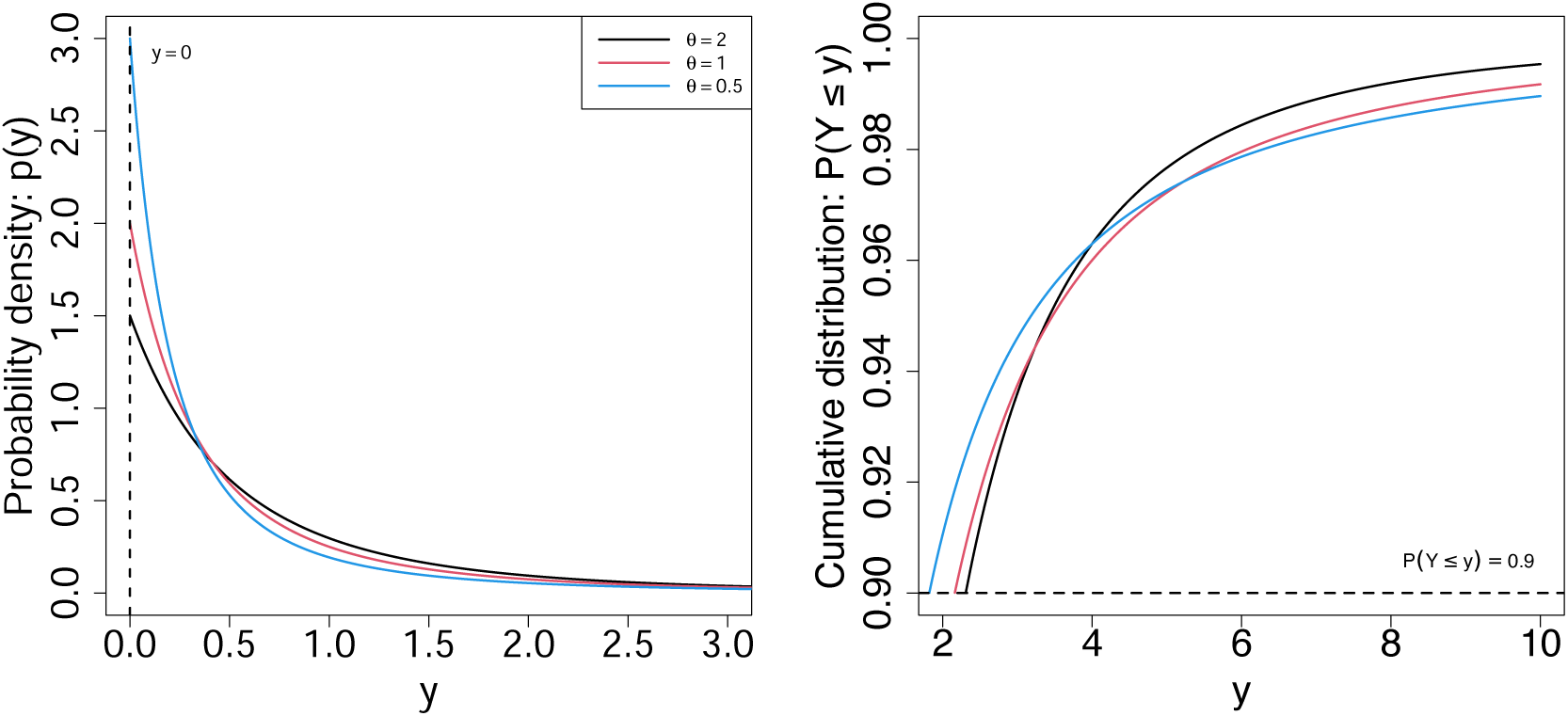
Lomax probability density and cumulative distribution functions under three different values of *θ*.

Recall the sparsity and right skewness of MTX abundance for UniRef90_A0A015QIN6 in Section 2.1. Even after filtering using MGX abundance, 24% of observations remain zero, with a 75th percentile of 0.9895 and values ranging from 0 to 34.9527. Therefore, the heavy-tailed nature of the model is well suited to MTX data, as a lighter-tailed model may fit the concentration near zero only at the cost of underestimating the right tail.

As a limiting property, the Lomax random variable converges to 0 in probability as *θ* → 0, whereas as *θ* → ∞, it converges in distribution to an exponential distribution with scale parameter *µ_ij_* (Proposition A.5, Supplementary Material). The former indicates a potential limitation under extreme sparsity. Although the tail becomes increasingly heavy, *P* (*Y_ij_ > y*) → 0 for any fixed *y >* 0 as *θ* → 0. Then, when extreme sparsity is present in the data, it may compromise the description of nonzero observations by excessively concentrating the Lomax distribution near zero, despite an increasingly heavy tail. This consideration motivates the complete model introduced in Section 3.3.

### 3.3 Complete Model Specification

This section completes the full model in two steps, building on our base framework. First, we replace the inverse-gamma mixing distribution with a two-component inverse-gamma mixture to address the limitation under extreme sparsity identified in Section 3.2. Second, we incorporate a left-censoring mechanism to account for zero-valued observations that may arise from latent abundances below the detection limit. With these components, the complete exponential scale mixture (ESM) model is defined and used for subsequent posterior inference and analyses.

#### 3.3.1 Two-Component Mixture Prior for the Latent Scale

As discussed in Section 3.2, the parameter *θ* controls both the concentration of probability near zero and the heaviness of the right tail, with smaller values corresponding to greater concentration near zero and heavier tails. Extremely sparse MTX data may lead to a fitted value of *θ* near zero, concentrating most of the probability in a small neighborhood of the origin and constraining the model’s ability to describe nonzero measurements. We therefore seek to decouple the concentration near zero from the behavior over positive abundances by generalizing the inverse-gamma mixing distribution to a two-component mixture. The two components have distinct parameters *θ*_1_ and *θ*_2_: a smaller value for the strong near-zero concentration and a larger value for appreciable density over the positive range. Specifically, we assume

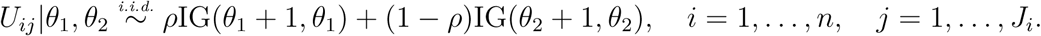

Marginalizing over *U_ij_*extends the marginal representation in (1) to a two-component Lomax distribution whose components share the same mean *µ_ij_* but differ in their precision parameters:

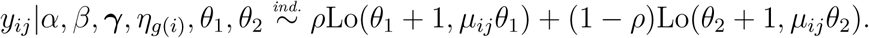

Consequently, this mixture formulation, with no change to the mean regression structure, reduces the reliance on a single *θ* to capture both near-zero concentration and right-tail behavior.

#### 3.3.2 Left-Censoring Mechanism for Zero Abundances

Zero measurements in MTX data may be the result of the underlying transcript abundance falling below the detection limit, for example, when the abundance is too low to be detected during preprocessing by a metatranscriptomic pipeline. Therefore, we treat a recorded zero as a censored realization of an underlying abundance *ỹ_ij_*. Specifically, we observe *y_ij_* = 0 if *ỹ_ij_ < C*, and *y_ij_* = *ỹ_ij_* otherwise, for a given threshold *C*. Let *F_ij_* denote the conditional cumulative distribution function of the underlying abundance given *λ_ij_* = *µ_ij_U_ij_*, such that *F_ij_*(*C*) = Pr(*Ỹ_ij_* < *C* | *λ_ij_*) = 1 − exp(−*C/λ_ij_*). Under this formulation, the contribution of the (*i, j*)th MTX observation to the likelihood changes from the exponential density exp(−*y_ij_/λ_ij_*)*/λ_ij_* to

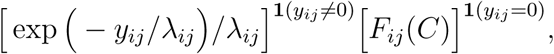

where **1**(*x* = *z*) denotes an indicator function that equals 1 if *x* = *z* and 0 otherwise. An interesting consequence of this formulation is that it assigns positive probability mass to an observed zero through *F_ij_*(*C*), rather than merely a positive density at zero as in the underlying exponential mixture, or equivalently Lomax, model. As a result, from a sampling perspective, the model can generate zero-valued MTX measurements with positive probability.

For posterior inference, we augment the model with an auxiliary latent variable *ỹ_ij_* satisfying 0 ≤ *ỹ_ij_ < C* for reported zero measurements, *y_ij_* = 0. We set *C* to the minimum nonzero MTX abundance observed across all genes, which serves as an empirical proxy for the minimum detectable abundance on the CPM scale. The augmentation allows posterior inference to remain computationally tractable, as described in Section F.2 of the Supplementary Material. Lastly, the hierarchical formulation of the complete model, which includes the two-component latent-scale specification and the left-censoring mechanism, is given in Section F.1 of the Supplementary Material.

## 4 Simulation Study

We conducted a simulation study using data from IBDMDB with a total of 1000 genes randomly selected from those analyzed in the case study in Section 5. Of the original 817 samples, those with missing metadata for any covariate were excluded, leaving 784 samples for the simulation study.

Then, we created a binary variable *s* by randomly dividing the samples into two equal groups, assigning one half to *s* = 1 and the other half to *s* = 0. A positive association was induced by multiplying the MTX measurements for samples with *s* = 1 by a signal strength of 5. Similarly, a negative association was induced by dividing the MTX measurements for samples with *s* = 1 by the same signal strength; equivalently, in terms of the group contrast, the MTX measurements for samples with *s* = 0 could instead be multiplied by this value. To select the genes to receive the injected signals, we randomly designated 250 genes (25% of the 1000 genes) as truly differentially expressed. Of these, 125 were assigned positive signals with *β* = log(5), and the remaining 125 were assigned negative signals with *β* = − log(5).

We repeated the random group assignment and signal injection 10 times to obtain 10 independent simulation replicates, and report the across-replicate mean of each performance measure in the results below. As a sensitivity analysis, we also repeated the simulation using 20 independent replicates. The results were qualitatively similar, and so the main conclusions remained unchanged. The corresponding analysis based on 20 replicates is shown in Supplementary Figure S6.

We conducted a comparative analysis with a representative existing method, the Gaussian model (GM) of Zhang et al. (2021), fitted using lmer() in the lmerTest package under four pseudocount choices: 0.01 (GM1), 0.001 (GM2), 0.0001 (GM3), and half the minimum nonzero abundance (GM4), to evaluate the relative performance of the proposed ESM model. For DE identification while accounting for multiple testing, we applied the Bayesian FDR procedure described in Section F.3 of the Supplementary Material to the ESM model and the Benjamini-Hochberg procedure to the Gaussian models.

Figure 4 summarizes the results in terms of differential expression detection, signal strength estimation, and goodness of fit. The ESM achieves a superior trade-off between the true positive rate (TPR) and empirical false discovery rate (FDR) – defined as the mean false discovery proportion (FDP) across simulation replicates – implying higher detection power at comparable realized FDR levels. The ESM model also provides more accurate estimates of signal strength, with smaller mean squared error (MSE). We evaluate signal strength using exp(|*β*^^^|), for which the true value is exp(|*β*|) = 5 for both positive and negative signals. The signal strength estimation is based on genes identified by the ESM model and all four Gaussian models within each replicate, with the resulting estimates pooled across genes and replicates.

**Figure 4:**
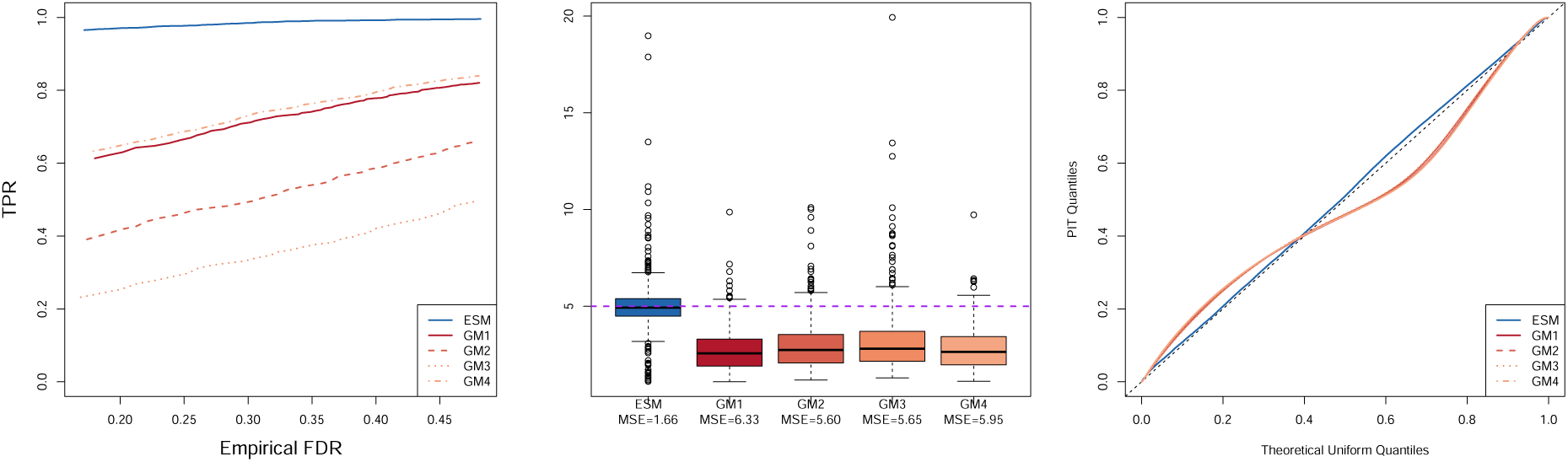
Simulation results comparing the ESM and GM models. From left to right: TPR versus FDR, boxplots of estimated signal strengths, and quantile–quantile (Q–Q) plots of probability integral transform (PIT) values against the standard uniform distribution. The dashed horizontal line in the middle panel indicates the true signal strength of 5, with mean squared errors (MSEs) reported below the model names. The diagonal line in the right panel represents perfect agreement with the standard uniform distribution.

We assessed goodness of fit using probability integral transform (PIT)-based diagnostics. An advantage of the PIT is that it maps observations onto a common probability scale. Thus, we can summarize the diagnostic results across genes and simulation iterations without consideration of standardization for differences in abundance scale. The common probability scale also permits direct comparison of the ESM fit to the original MTX abundances with the Gaussian fits to transformed abundances. For a well-fitting model, the PIT values are expected to follow a uniform distribution on (0, 1). PIT values from the ESM, pooled across genes and simulation iterations, closely follow the uniform reference distribution, whereas those from GM1–GM4 depart substantially from uniformity. This indicates markedly better goodness of fit under the ESM model and systematic lack of fit under the Gaussian models.

We further assessed distributional fit with replicated datasets (Table 2). Specifically, we examined Q25, Q50, Q75, and Q90 to assess different regions of the empirical abundance distribution. For the ESM model, replicated datasets were generated from the Lomax mixture distribution using model parameters drawn at each MCMC iteration. The selected quantiles were computed for each replicate, with pooling over MCMC iterations yielding posterior predictive distributions for Q25, Q50, Q75, and Q90, each summarized by a 95% interval.

**Table 2:** Predictive interval inclusion rates for selected distributional quantiles under the ESM and Gaussian models. Values represent the proportions of datasets, across 1000 genes and 10 simulation replicates, for which the observed quantile falls within the corresponding 95% predictive interval.

| Quantile | ESM | GM1 | GM2 | GM3 | GM4 |
|---|---|---|---|---|---|
| 25th percentile (Q25) | **0.959** | 0.272 | 0.253 | 0.256 | 0.265 |
| 50th percentile (Q50) | **0.929** | 0.038 | 0.010 | 0.005 | 0.044 |
| 75th percentile (Q75) | **0.931** | 0.196 | 0.149 | 0.130 | 0.201 |
| 90th percentile (Q90) | **0.870** | 0.242 | 0.218 | 0.199 | 0.264 |

For the Gaussian models estimated using lmer(), we used sim() from the arm package to obtain approximate posterior draws of the model parameters, analogous to the posterior draws used for the ESM model. Conditional on each parameter draw, a replicated dataset was generated from the corresponding Gaussian mixed model and transformed back to the original abundance scale, including subtraction of the corresponding pseudocount. The selected quantiles were computed for each replicate, with the resulting distributions summarized by 95% intervals.

We then calculated the proportion of gene–simulation combinations for which the observed Q25, Q50, Q75, and Q90 fell within their corresponding 95% interval estimates. Under the ESM model, these proportions were 0.959, 0.929, 0.931, and 0.870, respectively. In contrast, across the four Gaussian models, the corresponding proportions ranged from 0.253 to 0.272 for Q25, 0.005 to 0.044 for Q50, 0.130 to 0.201 for Q75, and 0.199 to 0.264 for Q90. Consequently, our proposed model reproduced the selected distributional features more accurately, and the GMs showed considerable discrepancies. This quantile-based assessment of replicated data complements the PIT diagnostic.

We additionally considered simulation settings in which the proportions of differentially expressed genes were 0.125 and 0.5, corresponding to 125 and 500 true DE genes, respectively, with the results reported in Supplementary Section C. Furthermore, Supplementary Figure S4 shows FDR calibration, in which the empirical FDR is plotted against the target FDR level. The GMs had clearly greater FDR inflation than the ESM model, particularly at low to moderate target FDR levels, indicating a greater false-discovery burden. FDR calibration under the ESM model was markedly better, even though some deviation from the target levels remained.

We also performed a sensitivity analysis of the left-censoring threshold for the ESM model. Following the strategy described in Section 3.3.2, we set the threshold to 0.00042, the minimum nonzero abundance across all genes and simulation replicates. We compared it with thresholds equal to one-half and one-tenth of the choice. The resulting TPR and FDP curves, together with the density estimates of the posterior means of *β*, were similar across the three specifications, indicating that the simulation results were robust to the choice of left-censoring threshold (Supplementary Figure S5).

## 5 Case Study

We conducted differential expression analysis with respect to dysbiosis using genes from the IBDMDB. We first applied the following screening criteria: genes were retained if at least two species were shared between the paired MTX and MGX data. We further screened the genes after MGX-based filtering to avoid perfect collinearity and model degeneracy. Specifically, dysbiosis and antibiotic use were required to be non-identical, with more than five observations in each category of both covariates. Lastly, the remaining genes were restricted to those with more than five nonzero MTX measurements in each dysbiosis category, alleviating the risk of estimated log-fold changes being driven by only a few nonzero observations.

The following analysis was based on 3000 genes randomly selected from those remaining after screening and the same 784 complete samples used in the simulation study, obtained after excluding observations with missing metadata for any covariate. The model variables {*s_i_, X_i_, g*(*i*)}, for *i* = 1*, . . .,* 784, are summarized as follows. For dysbiosis status, 123 observations were classified as “YES” and 661 as “NO”. Age had a mean of 28.18 years, with first and third quartiles of 14 and 42, respectively. For antibiotic use, 65 observations were “YES” and 719 were “NO”. The data set included 106 unique subjects, with *g*(*i*) ∈ {1*, . . .,* 106} denoting the subject identifier for sample *i*. An MGX-based filtering step discarded sample–taxon pairs (*i, j*) for which *D_ij_* = 0. We set the left-censoring threshold, *C*, to 0.000243529, the minimum nonzero MTX abundance observed across all genes.

Figure 5 compares the ESM model with GM1 as a representative Gaussian specification.

**Figure 5:**
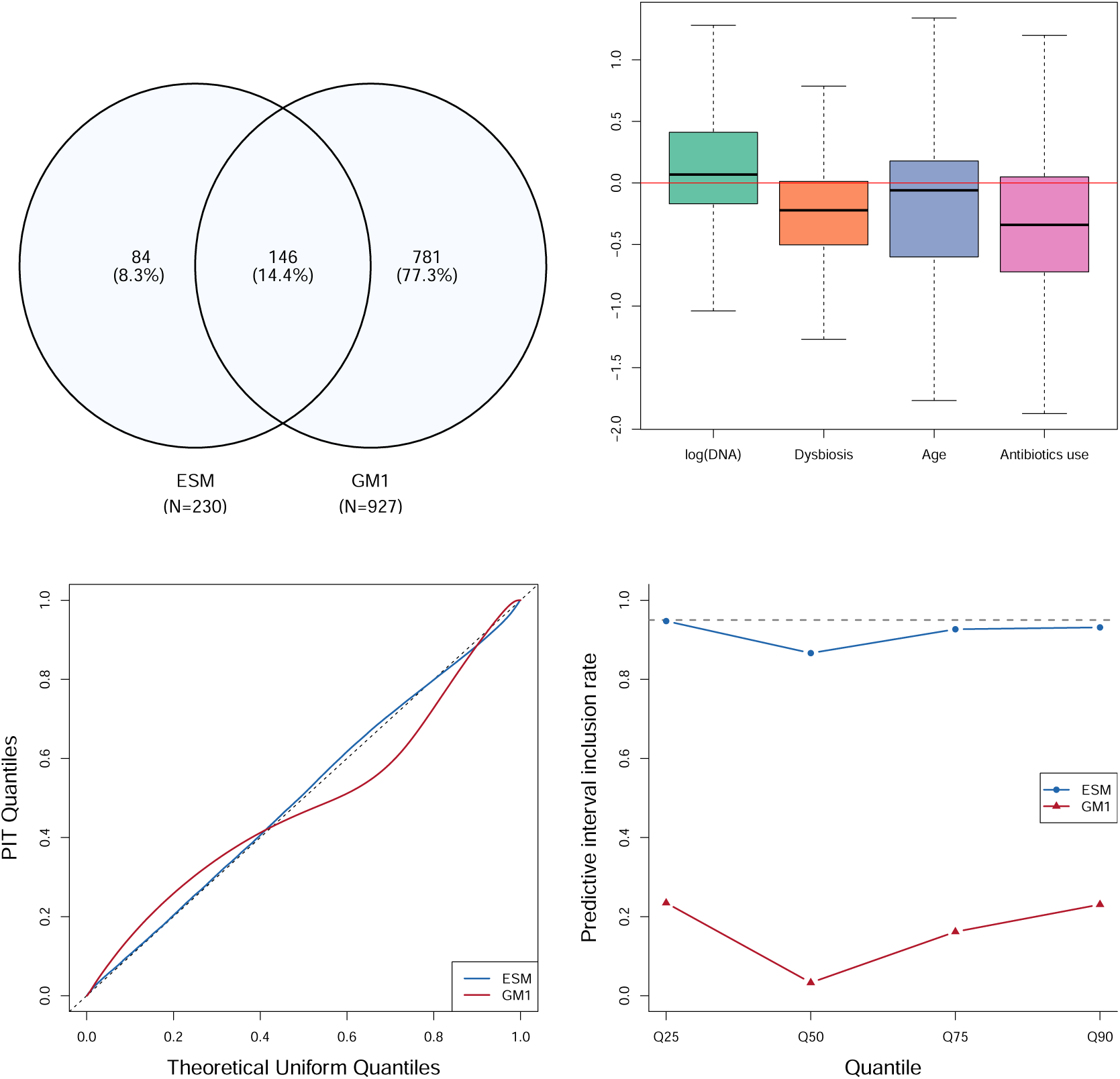
Comparison of the ESM and GM1 models in the IBDMDB analysis. Top left: overlap in genes identified as differentially expressed by the two models. Top right: boxplots of differences in estimated regression coefficients between the ESM and GM1 models for log(DNA), dysbiosis, age, and antibiotic use, with the horizontal line indicating zero difference (outliers omitted for display). Bottom left: Q-Q plots of PIT values against the standard uniform distribution. Bottom right: predictive interval inclusion rates for the 25th, 50th, 75th, and 90th percentiles, with the dashed horizontal line indicating the nominal 95% level.

Multiple testing adjustment was performed through the Bayesian FDR control described in Supplementary Section F.3 for the ESM model and the Benjamini–Hochberg method for GM1, with a target FDR level of *α* = 0.05. The ESM identified 230 genes as differentially expressed, whereas GM1 identified 927; 146 genes were common to both models, 84 were unique to the ESM, and 781 were unique to GM1. Such a large number of discoveries by GM1 should be interpreted cautiously given the severe FDR inflation of the Gaussian models observed in the simulation study. The top-right panel of the figure showing the difference in regression coefficient estimates between the two models indicates that their inferential discrepancies extended beyond the identification of DE genes. The distributional diagnostics strongly favored the ESM, with PIT values of the model closely following the standard uniform reference distribution, whereas those from GM1 showed substantial departures from uniformity. Consistently, the predictive interval inclusion rates under the ESM remained close to the nominal 95% level across the examined quantiles, while the corresponding rates under GM1 were substantially lower.

We also conducted a comparative analysis of the Gaussian model under different pseudocount choices and confirmed sensitivity to the pseudocount in both the identification of differentially expressed genes and the estimated effect sizes (see Figure S7 of the Supplementary Material).

For the genes exclusively identified by the ESM model, we performed functional annotation using the HUMAnN utility-mapping database, which allowed us to map the genes to KEGG orthologs (KOs) and their corresponding KEGG pathways, level-4 Enzyme Commission (EC) categories, eggNOG orthologous groups including COGs, and Pfam protein domains. Among the 84 exclusive genes, 65 received at least one functional annotation (see Table S5 of the Supplementary Material).

Then, we undertook a literature review for the 65 annotated genes to find prior evidence linking their functions to IBD and thus assess the potential biological relevance of these ESM-exclusive discoveries. UniRef90_A0A076ITY2 was annotated as a bacterial sialidase (K01186; EC 3.2.1.18), an enzymatic activity implicated in degradation of the intestinal mucus barrier in ulcerative colitis (Zhao et al. 2026). UniRef90_D4KXX9 was annotated as a peptidoglycan DL-endopeptidase (K21471), consistent with previous findings that gut microbial DL-endopeptidase abundance is reduced in Crohn’s disease and that experimental restoration of this activity can attenuate colitis through NOD2-mediated mechanisms (Gao et al. 2022). UniRef90_D4L453 was annotated as cysteine synthase A (CysK; K01738; EC 2.5.1.47), and *cysK* has been reported among microbial assimilatory sulfate-reduction genes with increased abundance in Crohn’s disease (Luo et al. 2024). In addition, several other ESM-exclusive genes, namely UniRef90_A0A3E4W374, UniRef90_D4KQG8, UniRef90_C7GCH8, UniRef90_D4KJ61, UniRef90_D4L2X3, were annotated to functions or pathways previously implicated in IBD, providing further biological context for these model-specific discoveries (Jangi et al. 2024, Bogdanova et al. 2026, Chen et al. 2026, Zhang et al. 2024, Luo et al. 2024).

To further examine functional relevance, we conducted Pfam enrichment analysis using the analyzed genes with available Pfam annotations as the background, with details provided in Section E of the Supplementary Material. Among the 230 genes identified as DE by the ESM, two Pfam domains associated with phosphoenolpyruvate (PEP)-utilizing enzymes, PF00391 and PF02896, were significantly enriched after Benjamini–Hochberg adjustment (adjusted *p* = 0.0366 for both). Each domain was annotated to five genes in the background set, four of which were identified as DE by the ESM, corresponding to a 9.6-fold enrichment. All four ESM-identified genes had positive estimated dysbiosis effects, with fold changes ranging from 1.58 to 2.63, whereas the remaining gene had a negligible estimated dysbiosis effect (*β*^^^ = −0.0004; PIP = 0.0824). In comparison, GM1 identified three of these four genes as differentially expressed, but neither domain was significantly enriched among its discoveries (fold enrichment = 1.79; adjusted *p* = 0.896). These findings suggest the potential functional relevance of the ESM discoveries involving microbial PEP-utilizing enzymes.

In summary, the ESM provides a more adequate distributional representation of the MTX data while identifying differentially expressed genes associated with biologically relevant microbial functions. In particular, several ESM-exclusive genes were annotated to functions that have previously been implicated in IBD. The enrichment analysis further identified two Pfam domains associated with PEP-utilizing enzymes, with consistently positive estimated dysbiosis effects across the four ESM-identified genes annotated with these domains.

## 6 Conclusion

Metatranscriptomic data are characterized by extreme sparsity; in the IBDMDB application, the proportion of zero-valued observations is as high as 0.9896 even at the 25th percentile across genes. We developed an exponential scale mixture (ESM) model for differential expression analysis that directly accommodates the highly concentrated and heavy-tailed abundance distributions encountered in such data, addressing both the distributional inadequacy and pseudocount sensitivity of a representative existing Gaussian model applied to transformed abundances.

The base exponential scale mixture framework is decomposed into a mean component and an observation-specific latent scale. Marginalizing over the scale parameter under an inverse-gamma mixing distribution yields a Lomax distribution whose mean is determined by the regression structure. Therefore, the regression coefficients are interpreted as log fold changes in mean abundance, while the parameter *θ* controls concentration near zero, right-tail behavior, and variance. The base framework’s limiting behavior reveals a potential difficulty under extreme sparsity. In this setting, the model may inadequately represent nonzero observations by excessively concentrating the Lomax distribution near zero. In particular, *P* (*Y_ij_ > y*) → 0 for any fixed *y >* 0 as *θ* → 0, even though the right tail becomes increasingly heavy. We tackle this limitation through a two-component Lomax mixture that accommodates extreme concentration near zero while maintaining appreciable density over the positive range. We also introduce a left-censoring mechanism for underlying transcript abundances that fall below the detection limit. From a generative perspective, the model can generate zero-valued MTX measurements by assigning positive probability mass to zero.

The simulation study demonstrates the inferential and distributional advantages of the complete ESM model. First, the proposed model achieved a better TPR–FDR trade-off, with higher detection power than the Gaussian models. It also yielded more accurate signal-strength estimates. Furthermore, the PIT diagnostic indicated better overall distributional fit. Quantitative checks using quantiles of replicated datasets showed that the ESM more closely reproduced features of the abundance distribution across multiple regions. The poorer distributional fit of the Gaussian models may propagate to downstream differential expression inference, which may partly explain their inferior DE performance.

The IBDMDB analysis provides empirical evidence for the practical utility of the proposed ESM model, while also revealing further differences from the Gaussian models in a real-data setting. The ESM and Gaussian models yielded markedly different results in DE gene identification, with GM1 producing substantially more discoveries. Given the severe FDR inflation observed for the Gaussian models in simulation, this larger number of discoveries should be interpreted cautiously. Distributional diagnostics favored the ESM, whereas the Gaussian specifications exhibited substantial lack of fit in qualitative and quantitative checks based on the PIT and replicated datasets, respectively. Beyond these methodological comparisons, the ESM identified dysbiosis-associated genes with functional annotations related to mucus-associated processes, peptidoglycan remodeling, and sulfur metabolism, all of which have previously been implicated in IBD. The enrichment analysis further found significant enrichment of two Pfam domains associated with PEP-utilizing enzymes, with consistently positive estimated dysbiosis effects across the four ESM-identified genes annotated with these domains, supporting the potential functional relevance of these discoveries.

Lastly, as a direction for further development, we will extend the proposed model, which analyzes genes independently and therefore does not model dependence across genes, to incorporate shared structure and borrow strength across genes. We will also consider evaluating the proposed approach in additional MTX cohorts to assess its generalizability.

