## Supplementary Material for "An Exponential Scale Mixture Model for Metatranscriptomics Data with Application to Inflammatory Bowel Disease"

Hyotae Kim and Li Ma

### A Proofs of Propositions

**Proposition A.1** (Marginalization). *Suppose* $U_{ij} \sim \mathrm{IG}(\theta+1, \theta)$ *and* $Y_{ij} \mid \mu_{ij}, U_{ij} \sim \mathrm{Exp}(\mu_{ij}U_{ij})$. *Then marginally*

$$Y_{ij} \sim \mathrm{Lo}(\theta+1, \mu_{ij}\theta).$$

*Proof.* The density marginalized over $U_{ij}$ is

$$\begin{aligned}
p(y_{ij}|\mu_{ij},\theta) &= \int_0^\infty (\mu_{ij}U_{ij})^{-1}\exp\Big(-y_{ij}(\mu_{ij}U_{ij})^{-1}\Big)\frac{\theta^{\theta+1}}{\Gamma(\theta+1)}U_{ij}^{-\theta-2}\exp(-\theta/U_{ij})dU_{ij} \\
&= \int_0^\infty \frac{\mu_{ij}^{-1}\theta^{\theta+1}}{\Gamma(\theta+1)}\frac{\Gamma(\theta+2)}{(\theta+y_{ij}\mu_{ij}^{-1})^{\theta+2}}\mathrm{IG}\Big(U_{ij}|(\theta+2),(\theta+y_{ij}\mu_{ij}^{-1})\Big)dU_{ij} \\
&= \frac{(\theta+1)}{\mu_{ij}\theta}\Big(1+\frac{y_{ij}}{\mu_{ij}\theta}\Big)^{-\big((\theta+1)+1\big)},
\end{aligned}$$

where $\mathrm{IG}(y|a,b)$ denotes the density of an inverse-gamma distribution $\mathrm{IG}(a,b)$. The resulting marginal density corresponds to that of the Lomax distribution $\mathrm{Lo}(\theta+1,\mu_{ij}\theta)$. □

**Proposition A.2** (Mean and variance). *For* $Y_{ij} \sim \mathrm{Lo}(\theta+1, \mu_{ij}\theta)$,

$$\mathrm{E}[Y_{ij}] = \mu_{ij}, \quad \theta > 0, \quad \mathrm{Var}(Y_{ij}) = \begin{cases} \mu_{ij}^2(\theta+1)/(\theta-1), & \theta > 1, \\ \infty, & 0 < \theta \le 1. \end{cases}$$

*Proof.* Under the hierarchical representation in (2), $Y_{ij} \mid U_{ij} \overset{ind.}{\sim} \mathrm{Exp}(\mu_{ij}U_{ij})$, where $U_{ij} \overset{i.i.d.}{\sim}$

$\mathrm{IG}(\theta+1,\theta)$. By the law of total expectation,

$$\begin{aligned}
\mathrm{E}[Y_{ij}] &= \mathrm{E}\Big\{\mathrm{E}[Y_{ij} \mid U_{ij}]\Big\} \\
&= \mu_{ij}\mathrm{E}[U_{ij}] \\
&= \mu_{ij}, \qquad \theta > 0.
\end{aligned}$$

Similarly, for $\theta > 1$, the second moment is

$$\begin{aligned}
\mathrm{E}[Y_{ij}^2] &= \mathrm{E}\Big\{\mathrm{E}[Y_{ij}^2 \mid U_{ij}]\Big\} \\
&= 2\mu_{ij}^2\mathrm{E}[U_{ij}^2] \\
&= \frac{2\mu_{ij}^2\theta}{\theta-1}.
\end{aligned}$$

Consequently,

$$\begin{aligned}
\mathrm{Var}(Y_{ij}) &= \mathrm{E}[Y_{ij}^2] - \Big\{\mathrm{E}[Y_{ij}]\Big\}^2 \\
&= \frac{2\mu_{ij}^2\theta}{\theta-1} - \mu_{ij}^2 \\
&= \frac{\mu_{ij}^2(\theta+1)}{\theta-1}.
\end{aligned}$$

For $0 < \theta \leq 1$, $\mathrm{E}[U_{ij}^2] = \infty$, and hence $\mathrm{Var}(Y_{ij}) = \infty$. □

**Proposition A.3** (Coefficient interpretability)**.** *Under the model* $\mu_{ij} = \exp\Big(\alpha\log(D_{ij}) + \beta s_i + \sum_{k=0}^{p}\gamma_k x_{ik} + \eta_{g(i)}\Big)$, *each coefficient* $\gamma_k$ *represents the log fold-change in the mean for a one-unit increase in covariate* $x_k$*, holding all other terms constant.*

*Proof.* Let $\mu_{ij}(x_{ik})$ denote the mean $\mu_{ij}$ evaluated at a specific value of $x_{ik}$, with all other covariates held fixed. Following Proposition A.2,

$$\mu_{ij}(x_{ik}) = \exp\Big(\alpha\log(D_{ij}) + \beta s_i + \gamma_{-k}^{\top}(X_i)_{-k} + \gamma_k x_{ik} + \eta_{g(i)}\Big),$$

where $(X_i)_{-k}$ denotes the covariate vector excluding $x_{ik}$, and $\gamma_{-k}$ is the corresponding coefficient vector excluding $\gamma_k$. Comparing two values $x_{ik}$ and $x'_{ik}$, we obtain

$$\frac{\mu_{ij}(x_{ik})}{\mu_{ij}(x'_{ik})} = \exp\Big(\gamma_k(x_{ik} - x'_{ik})\Big).$$

Because the model uses an exponential mean structure (i.e., a log link), multiplicative effects on the mean scale become additive on the log scale, making $\gamma_k$ interpretable as a log fold-change. $\square$

**Proposition A.4** (Tail behavior)**.** *For $Y_{ij} \sim \mathrm{Lo}(\theta+1, \mu_{ij}\theta)$, the distribution is regularly varying with tail index $(\theta+1)$ and is therefore heavy-tailed.*

*Proof.* The survival function is $\Pr(Y_{ij} > y) = \left(1 + \frac{y}{\mu_{ij}\theta}\right)^{-(\theta+1)} = y^{-(\theta+1)}L(y)$, where $L(y) = (1/y + 1/\mu_{ij}\theta)^{-(\theta+1)}$. Since $L(ty)/L(y) \to 1$ for all $t > 0$ as $y \to \infty$, $L(y)$ is slowly varying, and therefore the Lomax distribution is regularly varying with tail index $(\theta+1)$ $\square$

**Proposition A.5** (Limiting behavior in $\theta$)**.** *Let $Y_{ij}^{(\theta)} \sim \mathrm{Lo}(\theta+1, \mu_{ij}\theta)$.*

1. *As $\theta \to \infty$, $Y_{ij}^{(\theta)} \to \mathrm{Exp}(\mu_{ij})$ in distribution.*
2. *As $\theta \to 0^+$, $Y_{ij}^{(\theta)} \to 0$ in probability.*

*Proof.* (1) The cumulative distribution function of $Y_{ij}^{(\theta)}$ is

$$\Pr(Y_{ij}^{(\theta)} < a) = 1 - \left(1 + a/(\mu_{ij}\theta)\right)^{-(\theta+1)}.$$

Then

$$\lim_{\theta\to\infty} \Pr(Y_{ij}^{(\theta)} < a) = 1 - \left[\lim_{\theta\to\infty}\left(1 + a/(\mu_{ij}\theta)\right)^{\theta+1}\right]^{-1} = 1 - \left[\exp(a/\mu_{ij}) \times 1\right]^{-1} = 1 - \exp(-a/\mu_{ij}),$$

which is the cumulative distribution function of an exponential distribution with scale parameter $\mu_{ij}$. Hence, $Y_{ij}^{(\theta)} \to \mathrm{Exp}(\mu_{ij})$ in distribution.

(2) For every $\epsilon > 0$,

$$\Pr(Y_{ij}^{(\theta)} > \epsilon) = \left(1 + \epsilon/(\mu_{ij}\theta)\right)^{-(\theta+1)}.$$

As $\theta \to 0^+$, we have $(\theta+1) \to 1^+$ and $\epsilon/(\mu_{ij}\theta) \to \infty$, so $\Pr(Y_{ij}^{(\theta)} > \epsilon) \to 0$. By definition, $Y_{ij}^{(\theta)} \to 0$ in probability. $\square$

### B Sensitivity Analysis for Hyperparameter Choices

| | $\psi = 5$ | $\psi = 10$ | $\psi = 15$ | $\psi = 20$ |
|---|---|---|---|---|
| TPR/FPR/FDP | 0.83/0.03/0.06 | 0.81/0.03/0.05 | 0.80/0.03/0.06 | 0.82/0.03/0.05 |

Table S1: TPR, FPR, and FDP across values of $\psi$.

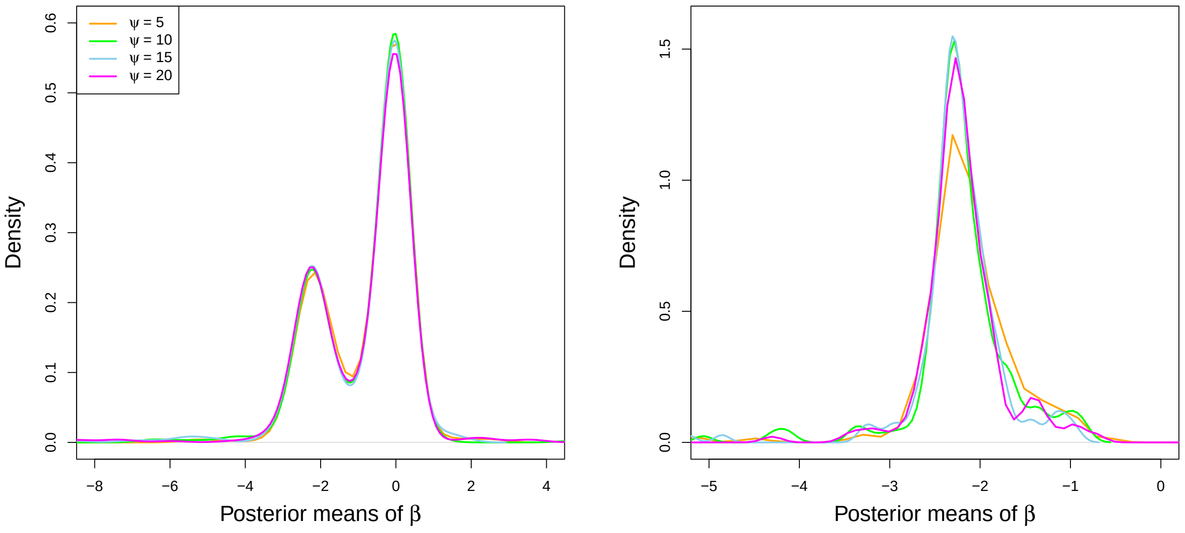


Figure S1: Densities (estimated using the `density` function in R) of the posterior means of $\beta$ for all 500 genes (left) and for genes identified as DE from models with different values of $\psi$ (right).

We conducted sensitivity analyses to assess the impact of the hyperparameters $\psi$ and $\xi$. As in the simulation study in the main paper (Section 4), we first generated IBDMDB-based synthetic data for the 500 selected genes. For each gene, we created a new grouping variable $s$, assigning a value of 1 to half of the zero MTX observations and half of the nonzero observations. Then, for a random subset of 200 (40%) genes, we induced a negative signal by multiplying MTX abundances by a factor of 10 for observations with $s = 0$.

We first considered four values of $\psi$ (5, 10, 15, and 20) and assessed sensitivity with respect

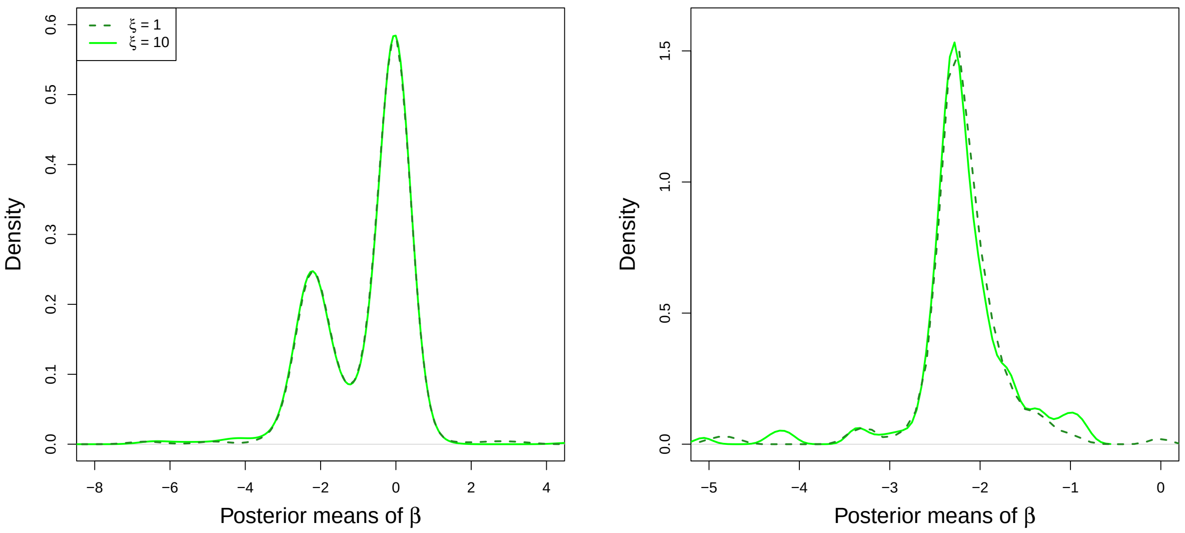


Figure S2: Densities of the posterior means of $\beta$ for all 500 genes (left) and for genes identified as DE (right) under $\xi = 1$ and 10.

to $\beta$, the parameter of primary interest in DE analysis. Figure S1 displays (estimated) densities of the posterior means of $\beta$ for all 500 genes and for subsets of genes identified as DE from models fitted with $\psi$ fixed at each value. The density based on all 500 genes indicates that the model is robust to the choice of $\psi$. The right panel for the DE subsets also shows robustness, although minor discrepancies are present; for example, the curve for $\psi = 5$ exhibits a slight deviation near the mode relative to the others. Table S1 reports the corresponding TPR, FPR, and FDP values, which are nearly identical across specifications, with differences on the order of $0.01 \sim 0.03$.

| | $\xi = 1$ | $\xi = 10$ |
|---|---|---|
| TPR/FPR/FDP | 0.79/0.03/0.05 | 0.81/0.03/0.05 |

Table S2: TPR, FPR, and FDP across values of $\xi$.

Similarly, we evaluated sensitivity to the choice of $\xi = 1, 10$ with respect to the key parameter $\beta$. Figure S2 shows densities of the posterior means of $\beta$, and Table S2 reports the corresponding TPR, FPR, and FDP, indicating robustness to the choice of $\xi$.

### C Additional Results for Simulation Study

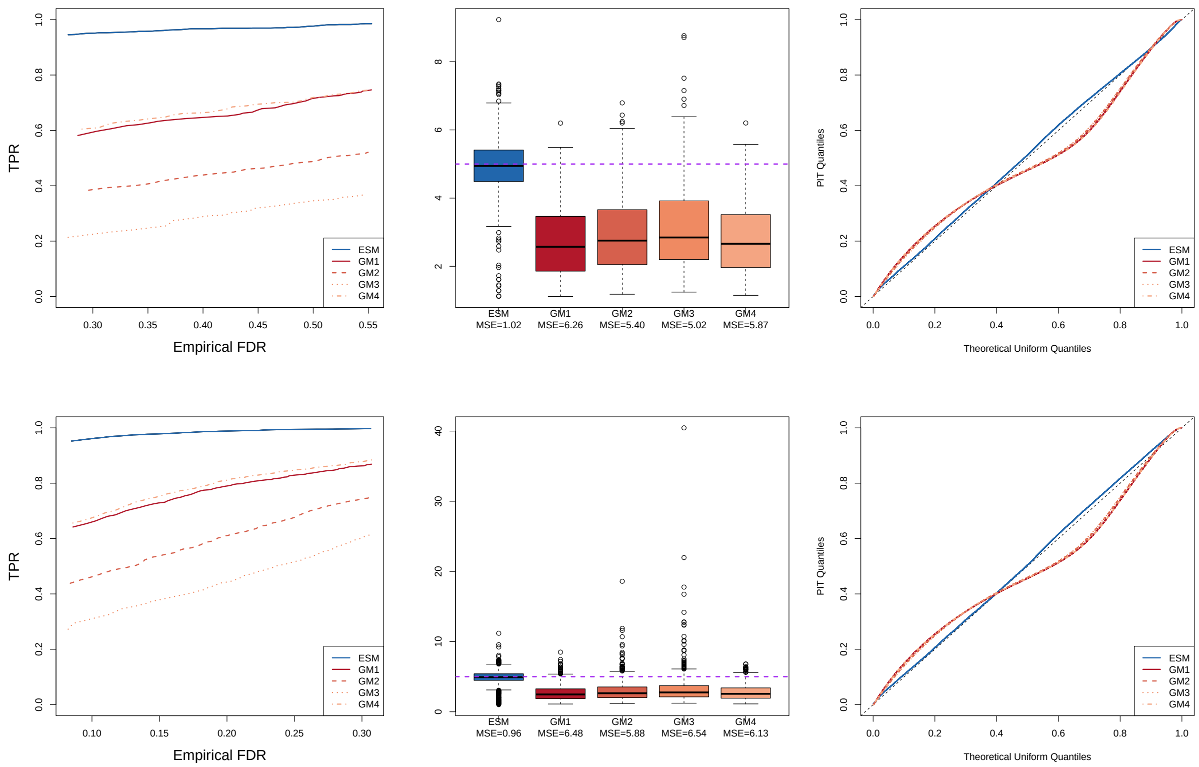


Figure S3: Simulation results comparing the ESM and GM models under two proportions of truly differentially expressed genes (0.125 and 0.5 in the top and bottom rows, respectively). From left to right: TPR versus FDR, boxplots of estimated signal strengths, and quantile–quantile (Q–Q) plots of probability integral transform (PIT) values against the standard uniform distribution. The dashed horizontal lines in the middle panels indicate the true signal strength of 5, with mean squared errors (MSEs) reported below the corresponding model names. The diagonal lines in the right panels represent perfect agreement with the standard uniform distribution.

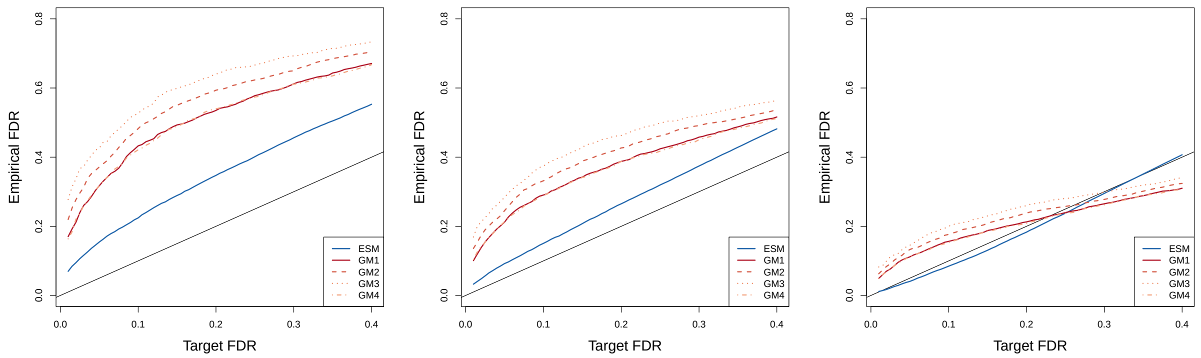


Figure S4: FDR calibration of the ESM and GM models under three proportions of truly differentially expressed genes (0.125, 0.25, and 0.5, from left to right). The empirical FDR is plotted against the target FDR level, with the black diagonal line indicating nominal FDR control.

| Quantile | ESM | GM1 | GM2 | GM3 | GM4 |
|---|---|---|---|---|---|
| 25th percentile (Q25) | **0.958** | 0.272 | 0.252 | 0.256 | 0.265 |
| 50th percentile (Q50) | **0.928** | 0.034 | 0.009 | 0.005 | 0.043 |
| 75th percentile (Q75) | **0.931** | 0.188 | 0.148 | 0.130 | 0.199 |
| 90th percentile (Q90) | **0.869** | 0.237 | 0.218 | 0.200 | 0.261 |

Table S3: Predictive interval inclusion rates for selected distributional quantiles under the ESM and Gaussian models with a proportion of truly differentially expressed genes of 0.125. Values represent the proportions of datasets, across 1000 genes and 10 simulation replicates, for which the observed quantile falls within the corresponding 95% predictive interval.

| Quantile | ESM | GM1 | GM2 | GM3 | GM4 |
|---|---|---|---|---|---|
| 25th percentile (Q25) | **0.960** | 0.271 | 0.253 | 0.255 | 0.264 |
| 50th percentile (Q50) | **0.932** | 0.048 | 0.012 | 0.006 | 0.049 |
| 75th percentile (Q75) | **0.931** | 0.211 | 0.154 | 0.132 | 0.206 |
| 90th percentile (Q90) | **0.872** | 0.251 | 0.222 | 0.203 | 0.269 |

Table S4: Predictive interval inclusion rates for selected distributional quantiles under the ESM and Gaussian models with a proportion of truly differentially expressed genes of 0.5. Values represent the proportions of datasets, across 1000 genes and 10 simulation replicates, for which the observed quantile falls within the corresponding 95% predictive interval.

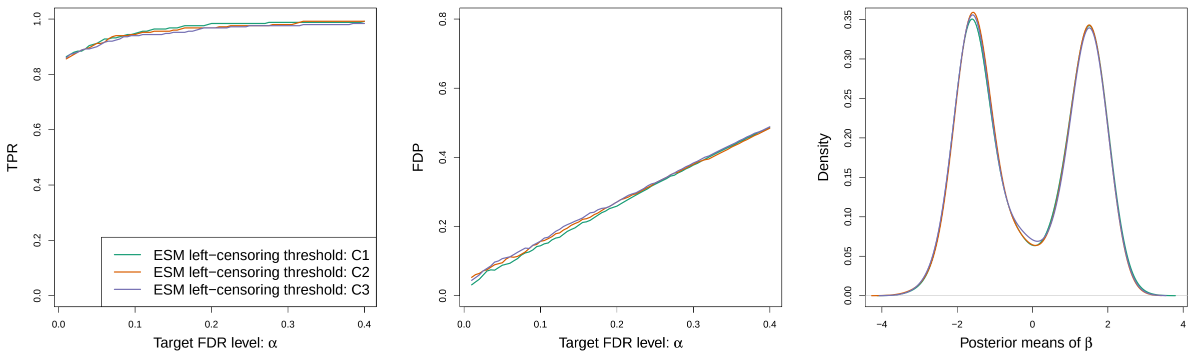


Figure S5: Sensitivity analysis with respect to the ESM model's left-censoring threshold. From left to right: TPR, FDP, and density estimates of the posterior means of $\beta$ for the identified genes under three left-censoring thresholds, $C_1 = 0.00042$, $C_2 = C_1/2$, and $C_3 = C_1/10$. TPR and FDP are shown across target FDR levels.

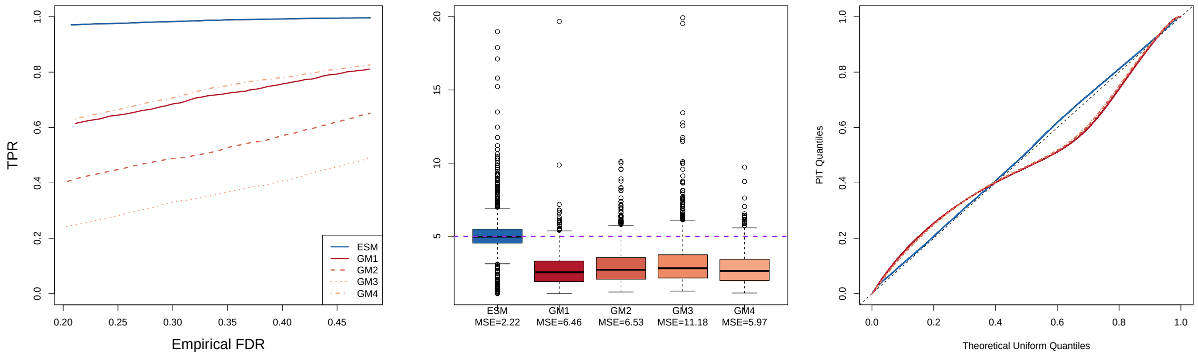


Figure S6: Simulation results comparing the ESM and GM models based on 20 simulation replicates. From left to right: TPR versus FDR, boxplots of estimated signal strengths, and quantile–quantile (Q–Q) plots of probability integral transform (PIT) values against the standard uniform distribution. The dashed horizontal line in the middle panel indicates the true signal strength of 5, with mean squared errors (MSEs) reported below the model names. The diagonal line in the right panel represents perfect agreement with the standard uniform distribution.

### D Additional Result for Case Study

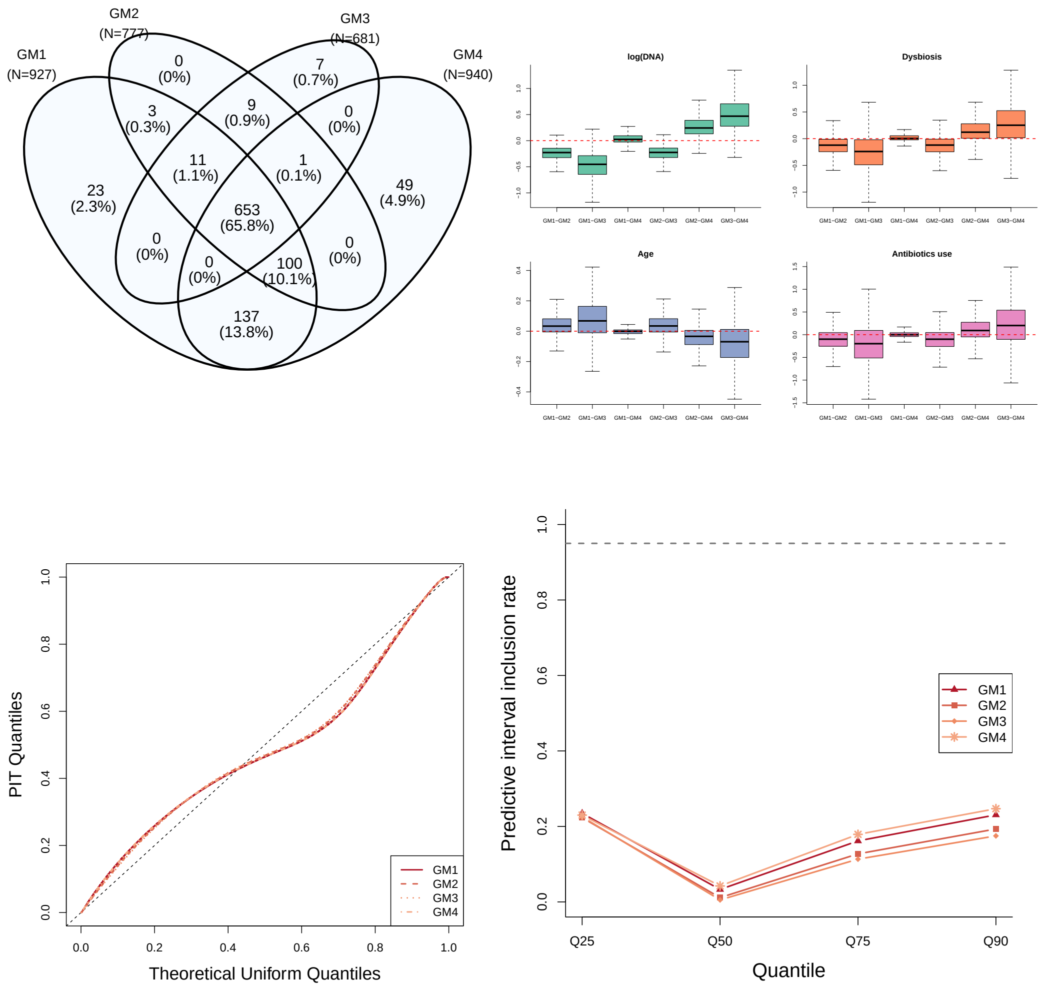


Figure S7: Comparison of the four Gaussian models under different pseudocount choices in the IBDMDB analysis. Top left: overlap in genes identified as differentially expressed by GM1-GM4. Top right: boxplots of pairwise differences in estimated regression coefficients among the four models for log(DNA), dysbiosis, age, and antibiotic use, with the horizontal line indicating zero difference (outliers omitted for display). Bottom left: Q-Q plots of PIT values against the standard uniform distribution, with the diagonal line representing perfect agreement. Bottom right: predictive interval inclusion rates for the 25th, 50th, 75th, and 90th percentiles, with the dashed horizontal line indicating the nominal 95% level.

Table S5: Functional annotations of the 84 gene families identified exclusively by the ESM. Gene families were annotated using UniRef90 functional descriptions, KEGG orthologs (KOs), level-4 Enzyme Commission (EC) categories, eggNOG/COG orthologous groups, Pfam protein domains, and KEGG pathways. A dash indicates that no corresponding annotation was available.

| **UniRef90** | **Gene name** | **KO** | **EC** | **eggNOG/COG** | **Pfam** | **KEGG pathway** |
|---|---|---|---|---|---|---|
| A0A015U8H2 | – | – | – | – | Immunity protein 17 | – |
| A0A076INQ7 | – | – | – | – | – | – |
| A0A076IRJ0 | Transcriptional regulator | – | – | – | MerR HTH family regulatory protein | – |
| A0A076ITY2 | Sialidase | sialidase-1 [EC:3.2.1.18] | – | – | BNR repeat-like domain; N-terminal domain of BNR-repeat neuraminidase | Other glycan degradation; Sphingolipid metabolism; Metabolic pathways; Lysosome biogenesis |
| A0A076IWK6 | – | – | – | – | – | – |
| A0A076J355 | Membrane protein | – | – | – | – | – |

*Continued on next page*

*Table S5 continued*

| UniRef90 | Gene name | KO | EC | eggNOG/COG | Pfam | KEGG pathway |
|---|---|---|---|---|---|---|
| A0A076J5N7 | 2-oxoglutarate oxidoreductase | 2-oxoglutarate/2-oxoacid ferredoxin oxidoreductase subunit beta [EC:1.2.7.3 1.2.7.11] | – | – | Thiamine pyrophosphate enzyme, C-terminal TPP binding domain | Glycolysis / Gluconeogenesis; Citrate cycle (TCA cycle); Pyruvate metabolism; Butanoate metabolism; Other carbon fixation pathways; Metabolic pathways; Biosynthesis of secondary metabolites; Microbial metabolism in diverse environments; Carbon metabolism; 2-Oxocarboxylic acid metabolism |
| A0A078QBF2 | Pilus formation N terminal region family protein | – | – | – | Pilus formation protein N terminal region | – |
| A0A078QTA4 | Glycosyl transferase group 2 family protein | – | – | – | Glycosyl transferase family group 2 | – |
| A0A078S3W1 | Type I restriction enzyme R Protein | – | Type I site-specific deoxyribonuclease | – | Type I restriction and modification enzyme - subunit R C terminal; Type I restriction enzyme R protein N terminus (HSDR_N) | – |
| A0A0K6BVS0 | – | – | – | – | – | – |

*Continued on next page*

*Table S5 continued*

| UniRef90 | Gene name | KO | EC | eggNOG/COG | Pfam | KEGG pathway |
|---|---|---|---|---|---|---|
| A0A0M6W9E2 | – | – | – | – | – | – |
| A0A0M6X1I1 | ABC-type multidrug transport system, ATPase component | – | – | – | ABC transporter | – |
| A0A0P0F5A8 | 8-amino-7-oxononanoate synthase | 8-amino-7-oxononanoate synthase [EC:2.3.1.47] | 8-amino-7-oxononanoate synthase | – | Aminotransferase class I and II | Biotin metabolism; Metabolic pathways; Biosynthesis of cofactors |
| A0A0P0G0U0 | ATP synthase subunit b | F-type H+-transporting ATPase subunit b | – | – | ATP synthase B/B' CF(0) | Oxidative phosphorylation; Photosynthesis; Metabolic pathways |
| A0A0P0M440 | Pyruvate,phosphate dikinase | – | Pyruvate, phosphate dikinase | – | PEP-utilising enzyme, mobile domain; PEP-utilising enzyme, TIM barrel domain; Pyruvate phosphate dikinase, PEP/pyruvate binding domain | – |
| A0A127SR39 | Erm23S_rRNA_methyltrans | – | – | – | Ribosomal RNA adenine dimethylase | – |
| A0A139K2B2 | – | – | – | – | – | – |
| A0A139K5W1 | Leucyl/phenylalanyl-tRNA–protein transferase | – | Lysine/arginine leucyltransferase | – | Leucyl/phenylalanyl-tRNA protein transferase | – |

*Continued on next page*

*Table S5 continued*

| UniRef90 | Gene name | KO | EC | eggNOG/COG | Pfam | KEGG pathway |
|---|---|---|---|---|---|---|
| A0A139K9A4 | Thioesterase family protein | – | – | – | Thioesterase superfamily | – |
| A0A139KJ68 | DNA gyrase subunit B | – | Transferred entry 5.6.2.2 | – | DNA gyrase B; DNA gyrase B subunit, carboxyl terminus; Histidine kinase-, DNA gyrase B-, and HSP90-like ATPase; Toprim domain | – |
| A0A139KTH9 | Sugar-binding domain protein | – | – | – | Bacterial regulatory proteins, lacI family; Periplasmic binding proteins and sugar binding domain of LacI family | – |
| A0A139T3K7 | – | – | – | – | – | – |
| A0A173TEK3 | Anaerobic benzoate catabolism transcriptional regulator | – | – | – | Helix-turn-helix | – |
| A0A173USY3 | Bicarbonate transport ATP-binding protein CmpD | – | – | – | ABC transporter | – |
| A0A174BVL2 | ABC transporter | – | – | – | ABC transporter | – |
| A0A174IM57 | Tetratricopeptide repeat | – | – | – | Tetratricopeptide repeat | – |

*Continued on next page*

*Table S5 continued*

| UniRef90 | Gene name | KO | EC | eggNOG/COG | Pfam | KEGG pathway |
|---|---|---|---|---|---|---|
| A0A174K2D1 | Biopolymer transport protein | – | – | – | Biopolymer transport protein ExbD/TolR | – |
| A0A174PDW8 | ABC transporter ATP-binding protein | – | – | – | ABC transporter | – |
| A0A1D3KEW1 | Protein YcfR | – | – | – | Protein of unknown function (DUF1471) | – |
| A0A1E3AYT6 | Rubrerythrin | – | – | – | Rubrerythrin | – |
| A0A1G8DLB9 | – | – | – | – | – | – |
| A0A1Q6JQG6 | AraC family transcriptional regulator | – | – | – | Helix-turn-helix domain | – |
| A0A1Y3Z8G4 | – | – | – | – | – | – |
| A0A374N817 | Cell envelope biogenesis protein OmpA | – | – | – | OmpA family; Outer membrane protein beta-barrel domain | – |
| A0A380Z4P9 | Peptidase S24-like./Bacteriophage CI repressor helix-turn-helix domain | – | – | – | Peptidase S24-like | – |
| A0A396E9A6 | – | – | – | – | – | – |
| A0A3D2LV15 | Histidinol-phosphatase | – | – | – | – | – |

*Continued on next page*

*Table S5 continued*

| UniRef90 | Gene name | KO | EC | eggNOG/COG | Pfam | KEGG pathway |
|---|---|---|---|---|---|---|
| A0A3D4Y3B6 | YihA family ribosome biogenesis GTP-binding protein | – | – | – | – | – |
| A0A3E4EC37 | DUF4318 domain-containing protein | – | – | – | – | – |
| A0A3E4GKA7 | – | – | – | – | – | – |
| A0A3E4W374 | Butyrate kinase | – | Butyrate kinase | – | – | – |
| A0A3E4WBZ6 | SPOR domain-containing protein | – | – | – | – | – |
| A0A3E5BD25 | Biopolymer transporter ExbD | – | – | – | – | – |
| A0A3E5F5K2 | – | – | – | – | – | – |
| A5ZGJ8 | Phosphorylase family | – | – | Uridine phosphorylase | Phosphorylase superfamily | – |
| A6L6M5 | – | – | – | – | Protein of unknown function (DUF3408) | – |
| A6NXN1 | – | – | – | – | – | – |
| A8S337 | – | – | – | Class II aldolase adducin | Class II Aldolase and Adducin N-terminal domain | – |

*Continued on next page*

*Table S5 continued*

| UniRef90 | Gene name | KO | EC | eggNOG/COG | Pfam | KEGG pathway |
| --- | --- | --- | --- | --- | --- | --- |
| B0NQA3 | Ferritin | – | Bacterial non-heme ferritin | Stores iron in a soluble, non-[...]on homeostasis (By similarity) | Ferritin-like domain | – |
| B0PDJ9 | – | – | – | conjugative transposon protein | Helix-turn-helix domain | – |
| B5CNT5 | Transposase | – | – | Pfam:Transposase_8 | – | – |
| B7UPI6 | Glucose-6-phosphate isomerase | glucose-6-phosphate isomerase [EC:5.3.1.9] | Glucose-6-phosphate isomerase | – | Phosphoglucose isomerase | Glycolysis / Gluconeogenesis; Pentose phosphate pathway; Starch and sucrose metabolism; Amino sugar and nucleotide sugar metabolism; Metabolic pathways; Biosynthesis of secondary metabolites; Microbial metabolism in diverse environments; Carbon metabolism; Biosynthesis of nucleotide sugars |
| C0FSP6 | ABC transporter, solute-binding protein | – | – | transporter activity | Bacterial extracellular solute-binding protein | – |
| C7GCH8 | Cobalamin biosynthesis protein CobD | – | – | – | CobD/Cbib protein | – |

*Continued on next page*

*Table S5 continued*

| UniRef90 | Gene name | KO | EC | eggNOG/COG | Pfam | KEGG pathway |
|---|---|---|---|---|---|---|
| D0TT49 | – | – | – | – | AMP-binding enzyme; AMP-binding enzyme C-terminal domain | – |
| D1JK98 | NAD dependent epimerase/dehydratase family protein | – | – | – | NAD dependent epimerase/dehydratase family | – |
| D4KJ61 | Phosphoribosylformyl-glycinamidine cyclo-ligase | phosphoribosylformyl-glycinamidine cyclo-ligase [EC:6.3.3.1] | Phosphoribosylformyl-glycinamidine cyclo-ligase | – | AIR synthase related protein, C-terminal domain; AIR synthase related protein, N-terminal domain | Purine metabolism; Metabolic pathways; Biosynthesis of secondary metabolites |
| D4KKR8 | Amino acid ABC transporter membrane protein, PAAT family (TC 3.A.1.3.-) | L-cystine transport system permease protein | – | – | Binding-protein-dependent transport system inner membrane component | ABC transporters |
| D4KQG8 | Flagellar hook-length control protein | flagellar hook-length control protein FliK | – | – | Flagellar hook-length control protein FliK | Flagellar assembly |
| D4KR03 | D-aminoacyl-tRNA deacylase | D-aminoacyl-tRNA deacylase [EC:3.1.1.96] | D-aminoacyl-tRNA deacylase | – | D-Tyr-tRNA(Tyr) deacylase | – |

*Continued on next page*

*Table S5 continued*

| UniRef90 | Gene name | KO | EC | eggNOG/COG | Pfam | KEGG pathway |
|---|---|---|---|---|---|---|
| D4KWE0 | Exopolyphosphatase-related proteins | bifunctional oligoribonuclease and PAP phosphatase NrnA [EC:3.1.3.7 3.1.13.3] | – | phosphoesterase RecJ domain protein | DHH family; DHHA1 domain | Sulfur metabolism; Metabolic pathways; Microbial metabolism in diverse environments |
| D4KX30 | Adenosyl cobinamide kinase/adenosyl cobinamide phosphate guanylyltransferase | adenosylcobinamide kinase / adenosylcobinamide-phosphate guanylyltransferase [EC:2.7.1.156 2.7.7.62] | Adenosylcobinamide kinase; Adenosylcobinamide-phosphate guanylyltransferase | Adenosylcobinamide kinase | Cobinamide kinase / cobinamide phosphate guanyltransferase | Porphyrin metabolism; Metabolic pathways; Biosynthesis of cofactors |
| D4KXX9 | Cell wall-associated hydrolases (Invasion-associated proteins) | peptidoglycan DL-endopeptidase CwlO [EC:3.4.-.-] | – | NLP P60 protein | Bacterial SH3 domain; NlpC/P60 family | – |
| D4KXZ4 | Tim44-like domain | – | – | – | Tim44-like domain; zinc-ribbon domain | – |
| D4KZW8 | Response regulator containing a CheY-like receiver domain and an HTH DNA-binding domain | – | – | Two component transcriptional regulator, LuxR family | Bacterial regulatory proteins, luxR family; Response regulator receiver domain | – |

*Continued on next page*

*Table S5 continued*

| UniRef90 | Gene name | KO | EC | eggNOG/COG | Pfam | KEGG pathway |
|---|---|---|---|---|---|---|
| D4L020 | Nucleotide-binding protein RO1_24910 | RNase adapter protein RapZ | – | Displays ATPase and GTPase activities (By similarity) | P-loop ATPase protein family | – |
| D4L196 | – | – | – | – | – | – |
| D4L2T8 | Shikimate kinase | shikimate kinase [EC:2.7.1.71] | Shikimate kinase | Catalyzes the specific phospho[...] a cosubstrate (By similarity) | Shikimate kinase | Phenylalanine, tyrosine and tryptophan biosynthesis; Metabolic pathways; Biosynthesis of secondary metabolites; Biosynthesis of amino acids |
| D4L2X3 | Sulfate adenylyltransferase subunit 1 | sulfate adenylyltransferase subunit 1 [EC:2.7.7.4] | Sulfate adenylyltransferase | may be the GTPase, regulating [...]ylase activity (By similarity) | Elongation factor Tu GTP binding domain | Purine metabolism; Monobactam biosynthesis; Selenocompound metabolism; Sulfur metabolism; Metabolic pathways; Biosynthesis of secondary metabolites; Microbial metabolism in diverse environments; Sulfur cycle |
| D4L3L3 | Endoribonuclease L-PSP, putative | – | – | Endoribonuclease LPSP | Endoribonuclease L-PSP | – |

*Continued on next page*

*Table S5 continued*

| UniRef90 | Gene name | KO | EC | eggNOG/COG | Pfam | KEGG pathway |
|---|---|---|---|---|---|---|
| D4L407 | Dinucleotide-utilizing enzymes involved in molybdopterin and thiamine biosynthesis family 2 | – | – | small protein activating enzyme activity | ThiF family | – |
| D4L453 | Cysteine synthase A | cysteine synthase [EC:2.5.1.47] | Cysteine synthase | cysteine biosynthetic process from serine | Pyridoxal-phosphate dependent enzyme | Cysteine and methionine metabolism; Sulfur metabolism; Metabolic pathways; Biosynthesis of secondary metabolites; Microbial metabolism in diverse environments; Carbon metabolism; Biosynthesis of amino acids; Sulfur cycle |
| D4LWJ1 | – | – | – | – | – | – |
| D5HH13 | Plasmid recombination enzyme | – | – | – | Plasmid recombination enzyme | – |
| D7IKR0 | Arabinose-proton symporter (Arabinose transporter) | – | – | – | Sugar (and other) transporter | – |
| D7IZV2 | – | – | – | – | Phosphodiester glycosidase | – |
| E1W5K2 | – | – | – | Membrane | Outer membrane efflux protein | – |

*Continued on next page*

*Table S5 continued*

| UniRef90 | Gene name | KO | EC | eggNOG/COG | Pfam | KEGG pathway |
|---|---|---|---|---|---|---|
| E4VYI3 | – | – | – | – | Domain of unknown function (DUF4369) | – |
| F3PFQ0 | Transglycosylase | – | – | penicillin-binding protein 1A | Penicillin binding protein transpeptidase domain; Transglycosylase | – |
| F3PRY0 | – | – | – | – | – | – |
| F7JMP3 | – | – | – | Resolvase | Domain of unknown function (DUF4368); Recombinase; Recombinase zinc beta ribbon domain; Resolvase, N terminal domain; Sigma-70 region 2 | – |
| G8FXJ6 | – | – | – | – | ABC-2 family transporter protein | – |
| I0PXY7 | – | – | – | – | – | – |

### E Pfam Enrichment Method

Pfam enrichment was assessed using an over-representation analysis based on the hypergeometric distribution. For each model, the foreground set consisted of genes identified as differentially expressed, and the background universe consisted of all analyzed genes with at least one Pfam annotation. Pfam gene sets were constructed from the corresponding gene-to-domain mappings, restricting attention to domains represented by at least 3 genes in the annotated background to avoid enrichment estimates driven by extremely small gene sets. For each Pfam domain, enrichment was evaluated using a one-sided hypergeometric test comparing the observed number of differentially expressed genes annotated to the domain with that expected under random selection from the annotated background. The expected count was calculated as $nM/N$, where $N$ is the number of genes in the annotated background, $n$ is the number of differentially expressed genes in that background, and $M$ is the number of background genes annotated to the domain. Fold enrichment was defined as the ratio of the observed to expected counts. Multiple testing was accounted for using the Benjamini-Hochberg procedure.

### F Posterior Inference

In Section F.1, we describe the hierarchical formulation of the proposed model, followed by details of its implementation in Section F.2. Section F.3 presents a Bayesian approach to differential expression inference with adjustment for multiple testing.

#### F.1 Hierarchical Specification

Below we present the full hierarchical representation of the complete model, including all prior distributions. The participant identifier for sample $i$ is denoted by $\text{PID}_i$, and

$\boldsymbol{y} = \{y_{ij} : i = 1, \ldots, n, \ \ j = 1, \ldots, J_i\}$. The model is given by

$$
\begin{aligned}
p(\boldsymbol{y}|\alpha, \beta, \boldsymbol{\gamma}, \{\eta_{g(i)}\}, \{U_{ij}\}) &\propto \prod_{i=1}^{n}\prod_{j=1}^{J_i} \Big[\exp\Big(-y_{ij}/(\mu_{ij}U_{ij})\Big)/(\mu_{ij}U_{ij})\Big]^{\mathbf{1}(y_{ij}\neq 0)} \Big[F_{ij}(C)\Big]^{\mathbf{1}(y_{ij}=0)} \\
\log(\mu_{ij}) &= \alpha \log(D_{ij}) + \beta s_i + \boldsymbol{\gamma}' X_i + \eta_{g(i)} \\
\alpha|\sigma^2 &\sim \mathrm{N}(0, \sigma^2) \\
\beta|\zeta, \tau^2 &\sim \begin{cases} \delta_0 & \text{if } \zeta = 0 \\ \mathrm{N}(0, \tau^2) & \text{if } \zeta = 1 \end{cases} \\
\zeta &\sim \mathrm{Bern}(\pi) \\
\tau &\sim \mathrm{C}^{+}(0, \psi) \Leftrightarrow \tau^2 \sim \mathrm{IG}(1/2, 1/b_{\tau^2}) \text{ and } b_{\tau^2} \sim \mathrm{IG}(1/2, 1/\psi^2) \\
\gamma_k|\upsilon_k^2 &\overset{i.i.d.}{\sim} \mathrm{N}(0, \upsilon_k^2), \quad k = 0, 1, \ldots, p \\
\eta_{g(i)}|\phi^2 &\overset{i.i.d.}{\sim} \mathrm{N}(0, \phi^2), \quad g(i) \in \{\mathrm{PID}_i : i = 1, \ldots, n\} \\
\sigma, \upsilon_k, \phi &\overset{i.i.d.}{\sim} \mathrm{C}^{+}(0, \psi) \\
U_{ij}|\theta_1, \theta_2, L_{ij} &\overset{i.i.d.}{\sim} \mathrm{IG}(\theta_{L_{ij}} + 1, \theta_{L_{ij}}), \quad i = 1, \ldots, n, \quad j = 1, \ldots, J_i \\
\Pr(L_{ij} = 1) = \rho &\quad \text{and} \quad \Pr(L_{ij} = 2) = 1 - \rho \\
\rho &\sim \mathrm{Unif}(0, 1) \\
\theta_1, \theta_2 &\overset{i.i.d.}{\sim} \mathrm{Exp}(\xi),
\end{aligned} \tag{S1}
$$

where $\mathrm{N}(\mu, \sigma^2)$ is the normal distribution with mean $\mu$ and variance $\sigma^2$, and $\mathrm{Bern}(\pi)$ refers to the Bernoulli distribution with mean $\pi$. $\mathrm{Unif}(0, 1)$ denotes an uniform distribution with support on the interval from 0 to 1. $\delta_0$ represents the Dirac delta function, which has zero density for all values except at 0. $\mathrm{C}^{+}(0, \psi)$ is the half-Cauchy distribution with its density given by $p(y) = 2/(\pi\psi)/(1 + y^2/\psi^2)$ for $y \geq 0$.

For hypothesis testing of $\beta$ in the DE analysis with respect to the variable $s_i$ (*dysbiosis*), we adopt a spike-and-slab prior. The formulation in (S1) is a hierarchical representation of the prior with the latent variable $\zeta$; after marginalizing over $\zeta$, it reduces to the standard form $(1-\pi)\delta_0 + \pi \mathrm{N}(0, \tau^2)$. To reflect a balanced prior belief regarding zero-nonzero $\beta$, we set $\pi = 0.5$.

All standard deviation parameters $\sigma$, $\upsilon_k$, and $\phi$, including $\tau$ (the standard deviation of the slab distribution), associated with the normal priors for the coefficients, are assigned half-Cauchy distributions. As a heavy-tailed prior, the half-Cauchy places more mass in the tails than exponentially tailed priors. Combined with its substantial mass near zero, it functions as an effective shrinkage prior. We set $\psi = 10$, chosen empirically based on sensitivity analyses over $\psi = 5, 10, 15$, and 20. Specifically, we examined how $\beta$, the key parameter of interest in DE analysis, varies across these choices. Results of the sensitivity analysis for a simulation example are provided in Section B, where the inference results, including TPR (true positive rate), FPR (false positive rate), and FDP (false discovery proportion), are consistent across settings, with small discrepancies on the order of $0.01 \sim 0.03$ in the rates. Lastly, the half-Cauchy distribution prior allows for a hierarchical representation via inverse-gamma distributions for each corresponding variance (as shown in (S1) for $\tau$), which facilitates posterior sampling by yielding closed-form full conditional distributions for the variance parameters and their hyperparameters.

The weight $\rho$ for the inverse-gamma mixture of $U_{ij}$ is assigned $\mathrm{Unif}(0, 1)$, a noninformative prior for a probability parameter. For the parameters $\theta_1$ and $\theta_2$, which do not admit conjugate priors, we adopt independent exponential priors. In addition to the prior being parsimonious, with only one hyperparameter, the exponential density is monotonically decreasing and therefore favors smaller nonnegative values, which may be reasonable given the sparsity characteristic of MTX data. We conducted a sensitivity analysis with $\xi = 1$ and 10 to assess the effect of $\xi$ on estimates for $\beta$, including TPR, FPR, and FDP, and found the results to be robust in this regard (see Section B), with a small discrepancy of

0.02 in TPR. We set $\xi = 10$ for all subsequent analyses.

#### F.2 Model Implementation

Posterior inference is conducted using a Gibbs sampler, where closed-form full conditional distributions are available for most model parameters. To obtain these closed-form expressions, slice sampling with data augmentation plays a key role. First, the cumulative probability $F_{ij}(C)$ can be represented as $F_{ij}(C) = \int_0^C \exp\Big(-t/(\mu_{ij}U_{ij})\Big)/(\mu_{ij}U_{ij})dt$, where $\mu_{ij} = \exp\{\alpha\log(D_{ij}) + \beta s_i + \boldsymbol{\gamma}'X_i + \eta_{g(i)}\}$. Then, by introducing the latent variables $\tilde{\boldsymbol{y}} = \{\tilde{y}_{ij} : y_{ij} = 0,\ i = 1,\dots,n,\ j = 1,\dots,J_i\}$, the augmented likelihood is given by:

$$p(\boldsymbol{y}, \tilde{\boldsymbol{y}}|\{\mu_{ij}\},\{U_{ij}\}) \propto \prod_{i=1}^{n}\prod_{j=1}^{J_i}\left[\frac{\exp(-y_{ij}/(\mu_{ij}U_{ij}))}{\mu_{ij}U_{ij}}\right]^{\mathbf{1}(y_{ij}\neq 0)}\left[\frac{\mathbf{1}(\tilde{y}_{ij} < C)\exp(-\tilde{y}_{ij}/(\mu_{ij}U_{ij}))}{\mu_{ij}U_{ij}}\right]^{\mathbf{1}(y_{ij}=0)}. \tag{S2}$$

With additional latent variables $d_{ij} > 0$, the exponential terms can be expressed as $\exp\Big(-\dot{y}_{ij}/(\mu_{ij}U_{ij})\Big) = \int_0^\infty \mathbf{1}\Big(d_{ij} < \exp\Big(-\dot{y}_{ij}(\mu_{ij}U_{ij})^{-1}\Big)\Big)dd_{ij}$, where $\dot{y}_{ij} = y_{ij}$ for observations with $y_{ij} \neq 0$ and $\dot{y}_{ij} = \tilde{y}_{ij}$ for those with $y_{ij} = 0$. Under the variable transformation $e_{ij} = -\log(d_{ij})$ with $\boldsymbol{e} = \{e_{ij} : i = 1,\dots,n,\ j = 1,\dots,J_i\}$, the augmented likelihood is written as

$$p(\boldsymbol{y}, \tilde{\boldsymbol{y}}, \boldsymbol{e}|\{\mu_{ij}\},\{U_{ij}\}) \propto \prod_{i=1}^{n}\prod_{j=1}^{J_i}\mathbf{1}\Big(e_{ij} > \dot{y}_{ij}(\mu_{ij}U_{ij})^{-1}\Big)\exp(-e_{ij})(\mu_{ij}U_{ij})^{-1}\Big[\mathbf{1}(\tilde{y}_{ij} < C)\Big]^{\mathbf{1}(y_{ij}=0)}. \tag{S3}$$

The full conditional distribution of $\tilde{y}_{ij}$ for observations with $y_{ij} = 0$, $i = 1,\dots,n$ and $j = 1,\dots,J_i$, is proportional to $\mathbf{1}(\tilde{y}_{ij} < C)\exp\Big(-\tilde{y}_{ij}/(\mu_{ij}U_{ij})\Big)$, as given in (S2). Consequently, $\tilde{y}_{ij}$ is sampled from a truncated exponential distribution, $\text{Exp}(\mu_{ij}U_{ij})$, with support restricted to $\tilde{y}_{ij} < C$.

The full conditional distribution for $e_{ij}$, $i = 1,\dots,n$, $j = 1,\dots,J_i$, is given by $p(e_{ij}|\mu_{ij},U_{ij},y_{ij},\tilde{y}_{ij}) \propto \Big(e_{ij} > \dot{y}_{ij}(\mu_{ij}U_{ij})^{-1}\Big)\exp(-e_{ij})$ from (S3). Therefore, $e_{ij}$ follows a truncated exponential distribution, Exp(1) with support $e_{ij} > \dot{y}_{ij}(\mu_{ij}U_{ij})^{-1}$. A posterior sample can be drawn by sampling from Exp(1) and adding $\dot{y}_{ij}(\mu_{ij}U_{ij})^{-1}$ to the result.

The full conditional for $\alpha$ is derived from (S3) as

$$\begin{aligned} p(\alpha|\beta, \boldsymbol{\gamma}, \{\eta_{g(i)}\}, \boldsymbol{y}, \tilde{\boldsymbol{y}}, \boldsymbol{e}) &\propto \prod_{i=1}^{n}\prod_{j=1}^{J_i}\Big[\mathbf{1}\Big(e_{ij} > \dot{y}_{ij}\exp(-\alpha\log(D_{ij}) - \beta s_i - \boldsymbol{\gamma}'X_i - \eta_{g(i)})U_{ij}^{-1}\Big) \\ &\quad \times \exp\Big(-\alpha\log(D_{ij})\Big)\Big]\exp\Big(-\alpha^2/(2\sigma^2)\Big) \\ &\propto \prod_{i=1}^{n}\prod_{j=1}^{J_i}\Big[\mathbf{1}\Big(-\log(e_{ij}U_{ij}/\dot{y}_{ij}) - \beta s_i - \boldsymbol{\gamma}'X_i - \eta_{g(i)} < \alpha\log(D_{ij})\Big)\Big] \\ &\quad \times \exp\Big(-\sum_{i=1}^{n}\sum_{j=1}^{J_i}\alpha\log(D_{ij}) - \alpha^2/(2\sigma^2)\Big). \end{aligned}$$

Therefore, posterior samples of $\alpha$ are drawn from a truncated normal distribution, $\mathrm{N}\Big(-\sigma^2\sum_{i=1}^{n}\sum_{j=1}^{J_i}\log(D_{ij}), \sigma^2\Big)$, with lower and upper truncation bounds given by $\max_{\{i,j:\log(D_{ij})>0\}}\Big[\frac{-\log(e_{ij}U_{ij}/\dot{y}_{ij}) - \beta s_i - \boldsymbol{\gamma}'X_i - \eta_{g(i)}}{\log(D_{ij})}\Big]$ and $\min_{\{i,j:\log(D_{ij})<0\}}\Big[\frac{-\log(e_{ij}U_{ij}/\dot{y}_{ij}) - \beta s_i - \boldsymbol{\gamma}'X_i - \eta_{g(i)}}{\log(D_{ij})}\Big]$, respectively.

Similarly, each coefficient $\gamma_k$ associated with covariate $X_{ik}$ is sampled from a truncated normal distribution, $\mathrm{N}\Big(-v_k^2\sum_{i=1}^{n}\sum_{j=1}^{J_i}X_{ik}, v_k^2\Big)$, with lower and upper truncation bounds given by $\max_{\{i,j:X_{ik}>0\}}\Big[\frac{-\log(e_{ij}U_{ij}/\dot{y}_{ij}) - \alpha\log(D_{ij}) - \beta s_i - \sum_{q\neq k}\gamma_q X_{iq} - \eta_{g(i)}}{X_{ik}}\Big]$ and $\min_{\{i,j:X_{ik}<0\}}\Big[\frac{-\log(e_{ij}U_{ij}/\dot{y}_{ij}) - \alpha\log(D_{ij}) - \beta s_i - \sum_{q\neq k}\gamma_q X_{iq} - \eta_{g(i)}}{X_{ik}}\Big]$, respectively.

The random effect $\eta_r$, for $r = 1, \ldots, R$, where $r$ indexes unique PIDs and $R$ denotes their total number, is also drawn from a truncated normal distribution, $\mathrm{N}\Big(-\phi^2\sum_{i:g(i)=r}\sum_{j=1}^{J_i}1, \phi^2\Big)$, with a lower truncation bound given by $\max_{\{i,j:g(i)=r\}}\Big[-\log(e_{ij}U_{ij}/\dot{y}_{ij}) - \alpha\log(D_{ij}) - \beta s_i - \boldsymbol{\gamma}'X_i\Big]$.

The coefficient $\beta$ for the DE analysis, assigned a spike-and-slab prior, is updated primarily according to $\zeta$: when $\zeta = 0$, $\beta = 0$; when $\zeta = 1$, $\beta$ follows a truncated normal distribution, $\mathrm{N}\Big(-\tau^2\sum_{i=1}^{n}\sum_{j=1}^{J_i}s_i, \tau^2\Big)$, with lower and upper bounds $\max_{\{i,j:s_i>0\}}\Big[\frac{-\log(e_{ij}U_{ij}/\dot{y}_{ij}) - \alpha\log(D_{ij}) - \boldsymbol{\gamma}'X_i - \eta_{g(i)}}{s_i}\Big]$ and $\min_{\{i,j:s_i<0\}}\Big[\frac{-\log(e_{ij}U_{ij}/\dot{y}_{ij}) - \alpha\log(D_{ij}) - \boldsymbol{\gamma}'X_i - \eta_{g(i)}}{s_i}\Big]$.

Regarding the update of $\zeta$, we first integrate out $\beta$ from the joint posterior distribution $p(\boldsymbol{y}, \dot{\boldsymbol{y}}, \boldsymbol{e}|\{\mu_{ij}\}, \{U_{ij}\})p(\beta)p(\zeta)$. Otherwise, $\zeta$ is deterministically defined by $\beta$: if $\beta = 0$, then $\zeta = 0$; if $\beta \neq 0$, then $\zeta = 1$. This leads to the full conditional distribution of $\zeta$ in the form $[(1-\pi)h_0]^{1(\zeta=0)}[\pi h_1]^{1(\zeta=1)}$, where $h_0 = \prod_{i=1}^n \prod_{j=1}^{J_i} \mathbf{1}\Big(e_{ij} > \dot{y}_{ij} \exp(-\alpha \log(D_{ij}) - \boldsymbol{\gamma}' X_i - \eta_{g(i)}) U_{ij}^{-1}\Big)$ and $h_1 = \exp\Big(\tau^2 (\sum_{i=1}^n \sum_{j=1}^{J_i} s_i)^2/2\Big) \int_{\text{LB}}^{\text{UB}} \text{N}(\beta| - \tau^2 \sum_{i=1}^n \sum_{j=1}^{J_i} s_i, \tau^2) d\beta$. The integration bounds are given by $\text{LB} = \max\limits_{\{i,j:s_i>0\}} \left[\frac{-\log(e_{ij}U_{ij}/\dot{y}_{ij}) - \alpha \log(D_{ij}) - \boldsymbol{\gamma}' X_i - \eta_{g(i)}}{s_i}\right]$ and $\text{UB} = \min\limits_{\{i,j:s_i<0\}} \left[\frac{-\log(e_{ij}U_{ij}/\dot{y}_{ij}) - \alpha \log(D_{ij}) - \boldsymbol{\gamma}' X_i - \eta_{g(i)}}{s_i}\right]$. Consequently, a posterior sample of $\zeta$ can be drawn from a Bernoulli distribution with success probability $\pi h_1 / \Big((1-\pi)h_0 + \pi h_1\Big)$.

Using the inverse-gamma representation of the half-Cauchy distribution, the variance parameters are sampled from updated inverse-gamma distributions. Specifically, for $(\sigma^2, b_{\sigma^2})$, the full conditionals are $\text{IG}(1/2 + 1/2, \alpha^2/2 + 1/b_{\sigma^2})$ and $\text{IG}(1/2 + 1/2, 1/\sigma^2 + 1/\psi^2)$, respectively. Similarly, for $(\upsilon_k^2, b_{\upsilon_k^2})$, the corresponding full conditionals are $\text{IG}(1/2 + 1/2, \gamma_k^2/2 + 1/b_{\upsilon_k^2})$ and $\text{IG}(1/2 + 1/2, 1/\upsilon_k^2 + 1/\psi^2)$. The parameter $\tau^2$ depends on the latent variable $\zeta$. When $\zeta = 1$ at a given MCMC iteration, $\tau^2$ is sampled from its inverse-gamma full conditional distribution, $\text{IG}(1/2 + 1/2, \beta^2/2 + 1/b_{\tau^2})$; otherwise, it is sampled from its prior distribution. The auxiliary parameter $b_{\tau^2}$ is sampled from $\text{IG}(1/2+1/2, 1/\tau^2+1/\psi^2)$. Finally, the variance parameter $\phi^2$ of the random effect is sampled from $\text{IG}(R/2 + 1/2, \sum_{r=1}^R \eta_r^2/2 + 1/b_{\phi^2})$, with $b_{\phi^2}$ sampled from $\text{IG}(1/2 + 1/2, 1/\phi^2 + 1/\psi^2)$.

The posterior sampling of $U_{ij}$ is based on the likelihood (S2) combined with its conjugate inverse-gamma prior, yielding $p(U_{ij}|\theta_1, \theta_2, L_{ij}, \mu_{ij}, y_{ij}, \tilde{y}_{ij}) \propto U_{ij}^{-1} \exp(-\dot{y}_{ij}\mu_{ij}^{-1}U_{ij}^{-1}) U_{ij}^{-\theta_{L_{ij}}-2} \exp(-\theta_{L_{ij}} U_{ij}^{-1})$. Consequently, $U_{ij}$ follows an updated inverse-gamma distribution, $\text{IG}(\theta_{L_{ij}} + 2, \dot{y}_{ij}\mu_{ij}^{-1} + \theta_{L_{ij}})$.

The mixture allocation indicator $L_{ij}$ is sampled from a discrete distribution with updated probabilities given by $\Pr(L_{ij} = 1) \propto \rho \text{IG}(U_{ij}|\theta_1 + 1, \theta_1)$ and $\Pr(L_{ij} = 2) \propto (1-\rho)\text{IG}(U_{ij}|\theta_2 + 1, \theta_2)$, where $\text{IG}(x|a, b)$ denotes the density function of an inverse gamma distribution $\text{IG}(a, b)$.

Parameter $\rho$ is updated using a beta distribution with parameters $n_1 + 1$ and $n_2 + 1$, where $n_1$ and $n_2$ denote the numbers of observations with $L_{ij} = 1$ and $L_{ij} = 2$, respectively.

The precision parameters $\theta_1$ and $\theta_2$ are the only variables that lack closed-form full conditional distributions; the Metropolis–Hastings algorithm is employed for their posterior sampling.

#### F.3 Posterior Inference for Differential Expression

Under the hierarchical formulation in (S1), $\zeta = 1$ indicates a nonzero $\beta$. Posterior inference for testing $\beta$ is conducted through the latent variable $\zeta$, specifically using the posterior inclusion probability (PIP), defined as the posterior mean of $\zeta$. The null hypothesis $\beta = 0$ is rejected if the PIP exceeds a prespecified threshold, such as 0.95 or 0.99, and is retained otherwise.

DE analysis typically involves testing many genes; multiple-testing adjustment is needed to control error rates such as the false discovery rate (FDR). Under our framework, applying such a fixed PIP threshold to each gene does not necessarily provide an efficient selection rule for a prespecified FDR level. Therefore, we adopt a Bayesian approach to FDR control: following the approach of Newton et al. (2004), we rank genes in ascending order of $1 - \text{PIP}$ and select the largest set of genes for which the average of $1 - \text{PIP}$ does not exceed the target FDR level. Formally, all genes $s \in S_m$ are declared differentially expressed if $\sum_{s \in S_m} (1 - \text{PIP}_s)/m \leq \alpha$, where $\alpha$ denotes the target FDR level, $S_m$ is the set of the first $m$ genes with the largest PIP values, and $m$ is the largest integer satisfying the inequality.
